# Transcription modulates chromatin dynamics and locus configuration sampling

**DOI:** 10.1101/2021.11.08.467739

**Authors:** Giada Forte, Adam Buckle, Shelagh Boyle, Davide Marenduzzo, Nick Gilbert, Chris A. Brackley

## Abstract

In living cells the 3D structure of gene loci is dynamic, but this is not revealed by 3C and FISH experiments in fixed samples, leaving a significant gap in our understanding. To overcome these limitations we applied the “highly predictive heteromorphic polymer” (HiP-HoP) model, validated by experiments, to determine chromatin fibre mobility at the *Pax6* locus in three mouse cell lines with different transcription states. While transcriptional activity minimally affects the movement of 40 kbp regions, we observed that the motion of smaller 1 kbp regions depends strongly on local disruption to chromatin fibre structure marked by H3K27 acetylation. This also significantly influenced locus configuration dynamics by modulating promoter-enhancer loops associated with protein bridging. Importantly these simulations indicate that chromatin dynamics are sufficiently fast to sample all possible conformations of loci within minutes, generating wide dynamic variability of gene loci structure within single cells. Experiments inhibiting transcription change chromatin fibre structure subtly, yet we predict they should substantially affect mobility. This combination of simulation and experimental validation provide a novel insight and mechanistic model to explain how transcriptional activity influences chromatin structure and gene dynamics.

---

The spatial organisation of the chromosome around a gene is thought to be intimately linked to its expression, providing an important mechanism for gene regulation. At short length scales (< 5000 bp), this mechanism might involve nucleosome repositioning and disruption of the chromatin fibre structure^1,2^, concomitantly modifying the accessibility of the DNA to proteins. At larger length scales (> 5000 bp), this could influence the way that chromatin forms loops bringing together promoters and their enhancers. Recent advances in microscopy^3,4^, and in next-generation sequencing methods such as chromosome-conformation-capture (3C) and its variants^5^, have revealed much about gene locus structure. More recently, the development of structural probes which operate at a single cell level have revealed a striking variability within a tissue or population of cells which are phenotypically homogeneous^3,4,6^. The compatibility of such cell-to-cell structural variability with a robust transcriptional programme and emerging phenotype is a remarkable phenomenon which still awaits a full explanation. The variability of gene loci was exemplified by our recent modelling work on the locus of *Pax6*, a highly-regulated and highly-conserved developmental gene. Our simulations showed that the chromatin interaction patterns revealed by CaptureC experiments could be generated by locus conformations which vary widely from cell to cell, and that the level of variation (validated using single cell microscopy using DNA FISH) also markedly depends on cell type and expression level^7^.

While experimental methods which probe chromosome organisation continue to improve, the majority of studies to date have focused on fixed cells. These only provide information on a “snapshot” in time, and an understanding of how locus conformations evolve dynamically remains largely elusive. For example, it is unclear whether the variation in 3D conformation observed across a population of cells is representative of the configurations adopted dynamically within a single cell. Or, does the chromatin in a single cell only visit, or sample, a small part of this “configuration space”, with the observed variability only arising when gathering all data from different cells? In other words, if one were to track a locus configuration in a live cell, would one observe wide variations, or a relatively static picture? The answer to this question may give insight into the wider issue of how such variability can still give tight control of expression and phenotypic homogeneity. Although live cell imaging and particle tracking methods have advanced significantly in recent years, there remain significant challenges. Super-resolution microscopy allows ever-increasing spatial resolution to be achieved, and high-throughput techniques such as Hi-D^8^ have been developed to monitor local diffusivity and flow of chromatin fibres *in vivo*. However, it remains challenging to reach high temporal resolution and to label multiple points of interest at the same time. In the present work we use biophysical modelling and computer simulations^9^ to study and predict the dynamics in the *Pax6* locus. This approach provides mechanistic insight as well as new hypotheses and testable predictions which we hope will stimulate further experiments.

Previously, we developed the “highly predictive heteromorphic polymer” (HiP-HoP) simulation framework to predict structural information on a gene locus, analogous to what can be observed in both single cell and population-level experiments. This model was applied to several gene loci, including those of *Pax6* in mouse, *SOX2* in human^7^, and (using an earlier version) the alpha and beta globin genes in mouse^10^. Here we evolved the HiP-HoP modelling approach to analyse the *dynamics* of the *Pax6* locus in three mouse tissue-derived cell lines which express the gene at different levels^11,12^. We first examined the dynamic properties of simulated chromatin regions at different length scales, finding that these vary substantially across the locus. The mobility of a given segment depends both on its biophysical properties (protein binding, local chromatin compaction), and on its surroundings, with local macromolecular crowding playing a pivotal role. We then studied the dynamics of the overall locus configuration, analysing the timescale over which promoter-enhancer interactions change. The results suggest that the locus can sample all its different configurations within minutes, strongly indicating that wide dynamic variation of locus structures would be observed within a single cell. Finally, we performed interventional experiments treating cells with either alpha amanitin to inhibit transcription or bleomycin to release topological strain; we made measurements in fixed cells, comparing these with simulations with the goal of inferring how this might affect the dynamics. Interestingly, we found that removing the proteins representing polymerase complexes from the simulation did *not* reproduce the changes observed in the alpha amanitin experiment; this led us to develop an alternative model scenario. This re-enforces the need to use biophysical modelling to understand chromatin structure and dynamics.

## Results

### Modified HiP-HoP framework to explore locus dynamics

We previously developed HiP-HoP, a polymer physics-based modelling framework^7,9,13^, to study the structure of the *Pax6* locus in three mouse cell lines, denoted *Pax6* OFF, ON and HIGH, to indicate the expression state of the gene. The model represents a chromosome region as a chain of beads and combines three key mechanisms which organise the locus (Fig. 1a). First, diffusing model proteins (representing complexes of RNA polymerase and transcription factors) can interact with the chromatin at specific binding sites; importantly these proteins are multivalent, and can bind to more than one chromatin site simultaneously forming molecular bridges^14,15^. Second, the loop extrusion mechanism is included^16–18^; this asserts that the cohesin complex can actively push chromatin into loops which grow until they are stabilised by CTCF proteins bound in a convergent orientation^19^. Finally, the model includes a “heteromorphic polymer” description of chromatin, which means that the biophysical properties, e.g. determined by disruptions of the chromatin fibre, can vary along its length^1,2,20^. For this, we discriminate between two chromatin states, one with a more compact thicker fibre, and one with a thinner less compact (more open, disrupted, or flexible) fibre^21^ (see Methods and Supplementary Information for details). In the present analysis we predominantly consider active chromatin regions, but this model could also be extended to account for repressed regions of the genome. An important feature is that proteins tend to come together into clusters; this is driven by a mechanism known as the “bridging induced attraction”^15,22^ and is a consequence of a protein’s ability to form molecular bridges. The clusters form around two or more protein binding sites on the chromatin and represent the foci or “phase-separated droplets” of transcription associated proteins observed in recent microscopy studies^23,24^. In HiP-HoP, the proteins continually switch back and forth between a chromatin binding and a non-binding state^25^ modelling post-translational modifications and resulting in clusters having realistic protein dynamics in terms of the exchange between the cluster and the soluble pool.

**Fig. 1.**
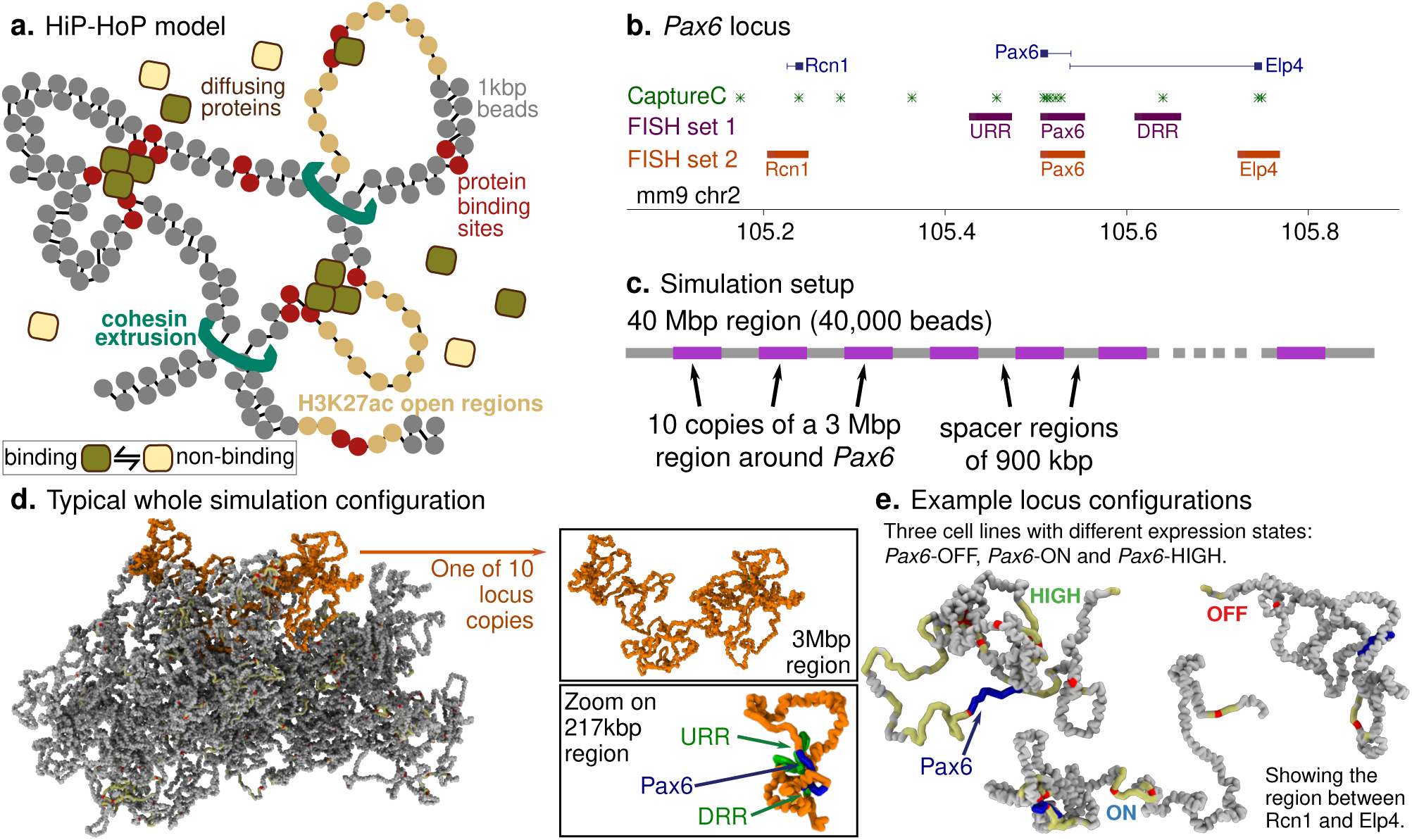
Model schematic and simulation set-up. **a**. In the HiP-HoP model a chromosome region is represented by a bead-and-spring polymer, where each bead represents 1 kbp of chromatin. The model includes diffusing protein complexes represented by spheres, loop extruders represented by additional springs, and a heteromorphic structure where some polymer regions have additional next-nearest neighbour springs which lead to local crumpling of the chain. Proteins stochastically switch back and forth between a binding and a non-binding state; when in the binding state they interact attractively with binding sites on the chromatin. Further details are given in Methods and Supplementary Information. **b**. A map of the mouse *Pax6* locus (mm9 genome build). Positions of CaptureC targets/viewpoints and two sets of FISH probes are indicated (see Supplementary Information for genomic coordinates). **c**. Schematic showing the simulation set up. Simulations of a 40 Mbp chromatin fibre (40,000 bead polymer) were performed. Ten copies of a 3 Mbp region around the *Pax6* locus were included on each fibre, allowing multiple results to be obtained from each simulation. Copies of the locus used input data from one of the three cell lines; in repeat simulations versions of the locus from different cell lines were in different positions. In this way each locus experienced a similar surrounding environment. **d**. Simulation snapshot showing a typical configuration of the 40 Mbp fibre. One of the 10 copies of the locus is shown in orange. In the top left this same copy of the locus is depicted with the rest of the fibre not visible. In the bottom left a zoom shows only the region immediately surrounding *Pax6* and two nearby regulatory regions (upstream and downstream regulatory regions, URR and DRR). **e**. Further simulation snapshots show example configurations of the locus (only the ∼ 500 kbp region between *Rcn1* and *Elp4* is shown) in each of three cell lines where *Pax6* is expressed at different levels. Chromatin regions with H3K27ac are shown in yellow, and ATAC-seq peaks (binding sites) in red. The *Pax6* gene body is shown in blue.

Three data sets are used as an input to the HiP-HoP model. First, DNA accessibility data (here ATAC-seq) is used to identify binding sites for the model proteins (we use a simplifying assumption that ATAC peaks coincide with binding sites for “active” proteins). Second, ChIP-seq data for acetylation of lysine 27 on histone H3 (H3K27ac) is used to identify regions which are in a more open chromatin state: this rational is based on several previous studies which suggest that chromatin possessing this mark has a disrupted structure^2,26^. And finally, ChIP-seq data for CTCF and cohesin (Rad21) are used to identify positions of loop-stabilising anchors (see Methods and Supplementary Information for all details).

In our previous study of the *Pax6* locus we used HiP-HoP to explore a population of simulated locus structures which also provided good predictions of experimental structural measures at both a population level (validated by CaptureC) and in single cells (validated by fluorescence *in situ* hybridization microscopy, FISH). CaptureC is a many-to-all 3C method which probes the genome-wide interactions of a set of selected target locations^27,28^. In this study our aim was to understand the *dynamics* of locus structures, providing new molecular insight; this necessitated significant changes to the simulation set up. We hypothesised that local variation in chromatin density might play a role in variation of its dynamics across the simulated region. Our previous simulations matched the *average* diffusion properties of the model chromatin to those measured experimentally, but since only a short region of the genome was simulated, spatial variations in chromatin density might not have been accurately represented. Consequently, in this work a much larger chromatin fragment (40 Mbp) was simulated to better match the overall density found *in vivo*. For computational efficiency, a region of 3 Mbp around the *Pax6* locus (chr2:104,000,000-107,000,000 mm9 genome build) was selected and concatemerized 10 times along the 40 Mbp fibre (i.e., one simulation is equivalent to taking measurements of the locus across 10 single cells, Figs. 1c,d). In a single simulation a mixture of loci from the three different cell lines was included, and in repeat simulations the positions of the loci in different cell lines were randomised (meaning that on average each copy of the locus was embedded in a similar chromatin context). In the simulations, time is evolved in discrete steps, with a simulation ‘time unit’ which can be mapped to a real time by comparison with experiments. Typically, each individual simulation was run through 1.5 million simulation time units (see Methods for more detail). Locus configurations from the first 700,000 time units were discarded as the simulations had not yet reached equilibrium (the configuration could still depend on the initial condition, see Supplementary Information); we then extracted 400 configurations from the remaining 800,000 time units for analysis (an example configuration from each of the three cell lines is shown in Fig. 1e).

To ensure that the simulations still gave good predictions of locus conformation with this new set up, the results were compared to both CaptureC data (Supplementary Fig. S1) and FISH (Supplementary Fig. S2). CaptureC targets were positioned at promoters and CTCF binding sites (including those within enhancer elements) across the locus (Fig. 1b, green stars). Three FISH probes were selected to cover the *Pax6* promoters and two previously identified enhancer regions, denoted the upstream and downstream regulatory regions (URR and DRR), and were used in 3-colour imaging experiments to obtain simultaneous measurements of probe separations in single cells (Fig. 1b, purple blocks). The conformations predicted by this new version of HiP-HoP showed a similar level of agreement with experimental data as previously^7^, but with the new simulation set up we could additionally measure chromatin interactions and separations as a function of time.

### Chromatin dynamics depend on transcription and fibre structure

To explore the dynamics around *Pax6* a simulation strategy was used that was analogous to undertaking live-cell imaging of specific regions of the locus, corresponding to the position of the *Pax6*, URR and DRR FISH probes (Fig. 2a). To achieve this the structure of the locus was simulated in all three cell lines (*Pax6* OFF, ON, HIGH). Results were obtained from 6 independent simulations, resulting in data equivalent to 20 single cells for each cell line. The mean squared displacement (MSD), as a measure of movement, was calculated as a function of lag time for each of the three regions covered by the FISH probes (Figs. 2b-d). In these simulations the length-scale of the chromatin (the diameter of the bead, denoted by *σ*), determined by comparing simulated and experimental FISH measurements, is 17.6 nm. The simulation time unit (denoted by *τ*) is approximately 2.07 ms, determined by comparison to previous motion tracking experiments^29^. This gives an approximate total duration for the analysed simulation of 28 minutes (see Methods and Supplementary Information for details of this mapping).

**Fig. 2.**
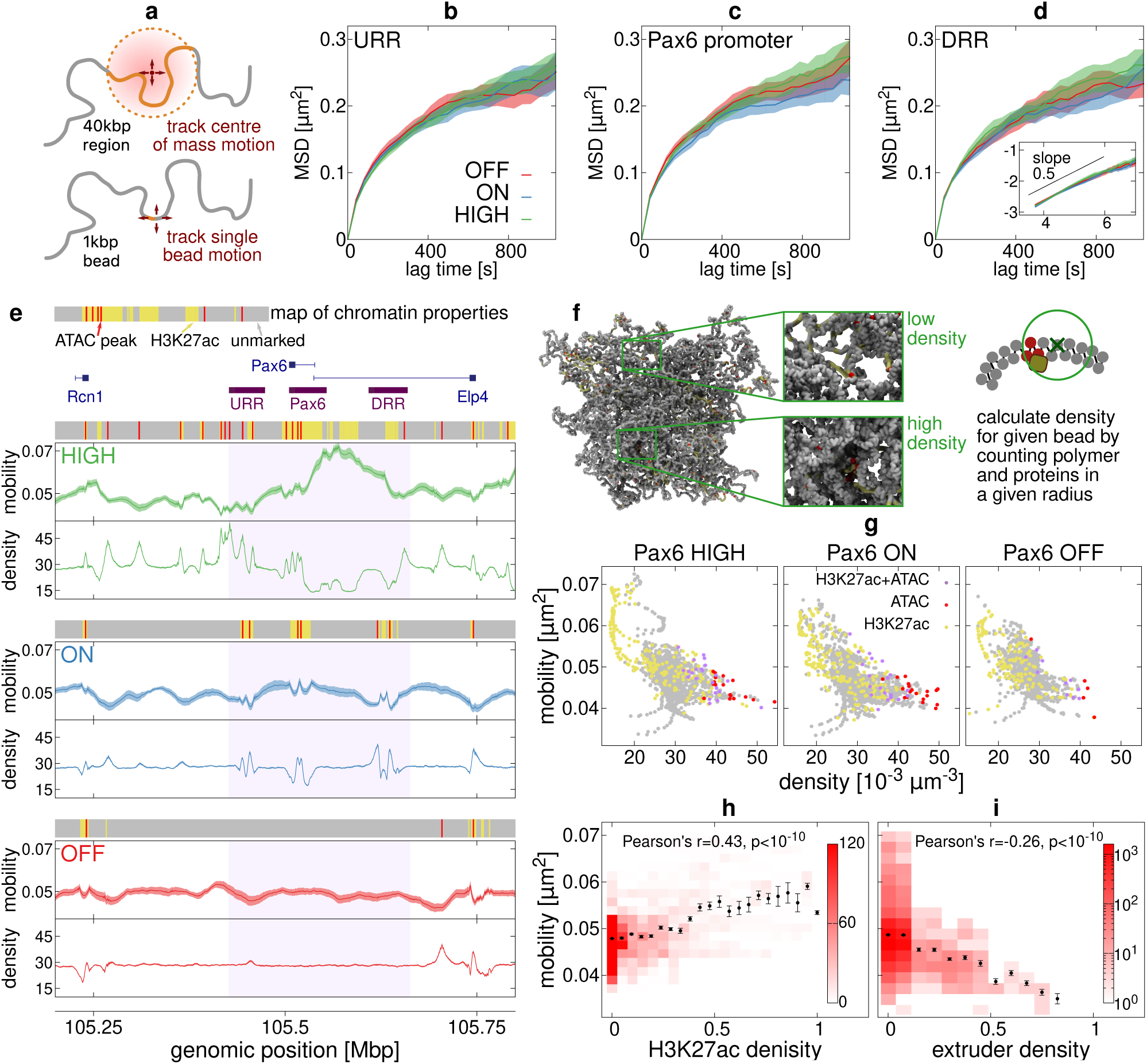
Dynamics of individual chromatin regions. **a**. To quantify the dynamics of a region covered by a FISH probe, the MSD of the centre of mass of the region covered by the probe was computed (top). Alternatively, the MSD of a single chromatin bead was computed (bottom). **b-d**. Plots showing the mean squared displacement (MSD) of simulated fluorescent probes positioned at the *Pax6* promoter and the two distal regulatory regions (URR and DRR). These cover the same regions as the FISH probes used in fixed cell experiments (probe set 1 in Fig. 1b). Simulation lengths and times were mapped to real units as detailed in Methods and Supplementary Information. All results are obtained from 20 independent simulations of the locus; results from three cell lines are shown (lines) and the shaded regions indicate the standard error in the mean. In panel c the inset shows log(MSD) *vs*. log(lag time), and the black line has a slope of 0.5. **e**. For each cell line, chromatin mobility is plotted as a function of position across the locus (units are ms^2^). Also shown is the mean local density (see panel f; units are 10^−3^ kbp *µ*m^−2^). The line indicates values for single polymer beads (each representing 1 kbp of chromatin); the shaded region indicates the standard error in the mean (for the density this is typically similar in size to the line width). Above each plot the coloured block shows the input data used for each cell line as indicated: yellow are H3K27ac regions, red are binding sites inferred from ATAC-seq peaks, other regions are grey. Gene positions are indicated above the plots. Purple blocks under the genes indicate the positions of the simulated FISH probes used to obtain the MSDs in panels b-d. **f**. The local density is determined for each chromatin bead by counting the number of proteins and polymer beads within a radius of 3*σ ≈* 53 nm. **g**. Scatter plots show the relationship between mobility and local density; each point represents a single polymer bead, and data from the whole 3 Mbp simulated region are included. Point colour indicates the type of bead (as determined from the input data). **h**. Plot showing the relationship between mobility and H3K27ac coverage. Values for each quantity are calculated for a 20 kbp window around each 1 kbp bead (the mean value of the mobility within the window, and the fraction of the 20 kbp region covered by the H3K27ac mark are used). To obtain the colour map, data are binned on both quantities, the darker the colour the more 20 kbp windows belong to the bin. To obtain points, data are binned according to the H3K27ac level only, and the mean mobility is calculated for each bin. Error bars show the standard error in the mean. The Pearson correlation coefficient and *p*-value are indicated. **i**. Plot showing the relationship between mobility and extruder occupancy. Extruder occupancy for a given bead is measured as the fraction of time which it is bound by an extruder, and we use the mobility for each bead. Again, the colour map is obtained by binning on both quantities, the points are obtained from binning on extruder occupancy with the mean mobility shown for each bin, and the Pearson correlation coefficient is indicated.

The MSD curves for different probes, and for a given probe in different cell lines look highly similar, indicating that the mobility of these regions is under similar constraints. For a freely diffusing object, the MSD grows linearly with time, but as expected for motion of a polymer segment, the MSDs here grew more slowly^30^ (inset Fig. 2d). Some slight differences between the cell lines can be seen at large times, indicating that the MSD grows more slowly with time for the *Pax6* ON cells compared to the others (particularly for the *Pax6* promoter probe and the DRR probe). Considering the input data which determines the chromatin properties, the *Pax6* ON cells have more protein binding sites (ATAC-seq peaks) than the other two cell lines, suggesting that interaction with proteins (stabilising chromatin loops) may lead to a local reduction in the mobility.

For each cell line the mobility *M* of the chromatin fibre (defined as the mean squared displacement MSD(Δ*t*) after a fixed lag time Δ*t* = 10^4^ *τ ≈* 20.7 s) was also calculated for each 1 kb bead to obtain a “mobility profile” over the locus (Fig. 2e). Focusing on the region immediately around *Pax6* and its regulatory elements (purple shaded region), there were large differences both across the locus, and between the cell lines. The most striking difference was around the DRR, which in *Pax6* HIGH cells shows a mobility around 1.5 times larger than in ON and OFF cells. In general, mobility tended to be higher in regions possessing H3K27ac (marked as yellow blocks above tracks), but lower at ATAC-seq peaks (marked as red bars). This is in stark contrast to the MSD measurements of the FISH probes where little difference was seen between probes or cell lines. This striking difference shows that dynamic measurements are highly sensitive to the way in which the chromatin is probed: the large local variation in mobility of different 1 kbp beads is largely obscured when analysing the ∼ 40 kbp simulated FISH probes, suggesting it is dependent on the behaviour of chromatin over small length scales. Examination of further simulated probes of different sizes revealed that both the observed mean and standard deviation of mobilities across the locus *decreased* with probe size (Supplementary Fig. 3c).

The diffusive motion of a chromatin segment is dependent on the spatial variation of chromatin and protein density which determines the effective local viscosity. This property is captured in our model by measuring the macromolecular density around each polymer bead (Fig. 2f). As predicted, lower local density is observed in regions which are more mobile (Fig. 2e) and quantitatively there was a significant negative correlation (Pearson correlation between − 0.5 and − 0.6, *p* < 10^−10^; see also Fig. 2g). It is also clear that the local density is enriched at model protein binding sites which can be visualised by colour coding the points in Fig. 2g. Beads located at ATAC-seq sites (red) had a 5.15% lower mobility on average than other beads. Reducing the number of model proteins in the simulations led to a reduction in local densities and an increase in mobility at binding sites (Fig S3), confirming that it is protein binding and bridge formation which impairs their motion. In contrast, H3K27ac marked beads (yellow in Fig. 2g) showed a 7.13% higher mobility on average than other beads. This is consistent with the more compact (non-H3K27ac) regions being more constrained within the fibre than the disrupted (H3K27ac) regions.

Increased mobility has previously been associated with increased levels of histone acetylation in live-cell imaging experiments^31^; consistently, there was a positive correlation between the mobility of 20 kbp windows and H3K27ac density in the simulations (Fig. 2h; Pearson’s *r* = 0.43, *p* < 10^−10^). To examine the effect of loop extrusion on locus dynamics, the extrusion probability *ϕ*_*e*_ for each bead was determined by measuring the fraction of time it was in the vicinity of an extruder. In the HiP-HoP framework extrusion probability is determined only by the positions and relative occupancy of CTCF sites and shows peaks at high CTCF occupancy sites (Supplementary Fig. S4). Extruders affected locus dynamics with a relatively weak correlation between extrusion probability and mobility (Fig. 2i).

A concern from this analysis is that the dynamics of a given chromatin bead is simply a reflection of the properties with which it is endowed in the simulation scheme. To test this the mobility of beads with no properties were analysed, and a broad range of mobility values were observed (Supplementary Fig. S5). This indicates that the dynamics of a given bead emerge both from its own properties (binding/non-binding open/compact) *and* from its local environment, which is dependent on the whole pattern of binding sites and chromatin states along the fibre^32^.

### Chromatin dynamics show positive correlation with interaction locality

We hypothesised that the chromatin properties which give rise to different dynamical behaviour might lead to differences in chromatin interactions, e.g., promoter-enhancer looping. A common measurement extracted from 3C-based experiments is the ratio between the amount of local and long ranged chromatin interactions at a given site^33^. Although this simulation setup limits the interpretation of long ranged interactions (many copies of a 3 Mbp locus are embedded in a larger chromatin fragment), a measure of “localness” of interactions can be determined in each cell type. This was parameterised as the ratio between the number of interactions with regions within 100 kbp of the target, and the number of interactions with regions further than 100 kbp (Fig. 3a, see Supplementary Information for details). 100 kbp was chosen as a threshold since it is slightly smaller than the expected mean promoter-enhancer loop size, so that cis-regulatory loops can be counted as long-ranged.

**Fig. 3.**
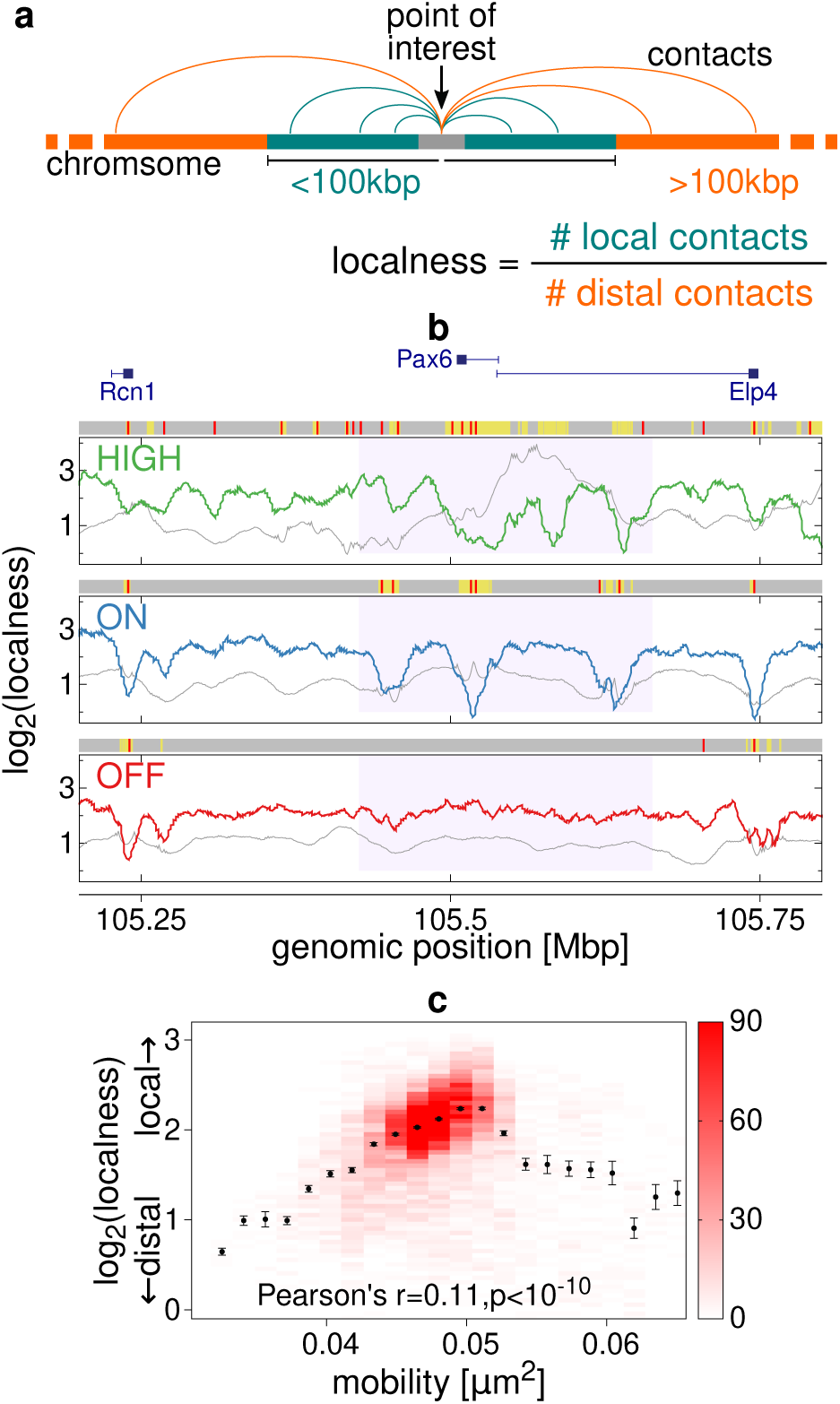
Dynamics are correlated with interaction locality. **a**. Localness is defined for a given chromatin bead as the number of interactions with regions closer than 100 kbp genomically, divided by the number of interactions with regions further away than 100 kbp. An interaction was defined as two chromatin beads being closer together than 3.5*σ ≈* 62 nm. **b**. Plots showing how localness of interactions varies across the locus in each cell type in simulations (coloured lines; a value is shown for each 1 kbp chromatin bead). log_2_ values are shown. For comparison, grey lines show the mobility as in Fig. 2e (a different vertical scale is used). Above each plot grey, yellow, and red bars indicate chromatin regions which were unmarked, H3K27ac marked (open chromatin) or ATAC-seq peaks (binding sites) in simulations. **c**. Plot showing the relationship between mobility and localness of interactions. To obtain the colour map, each chromatin bead was put into a bin depending on its localness and mobility; the darker the colour, the more beads belonging to the bin. Points were obtained by binning each chromatin bead only according to the mobility; the mean and standard error in the mean for each bin are shown. The Pearson correlation coefficient and *p*-value are indicated.

Measurement of localness for each 1 kbp bead across the simulated region (Fig. 3b) showed a pronounced reduction at protein binding sites (ATAC-seq peaks, red bars). This is consistent with these sites being involved in bridging interactions with distant (> 100 kbp) regions, leading to the creation of active protein clusters through the bridging-induced attraction^15,22^. Some, but not all, H3K27ac marked regions (open chromatin, yellow shading) also displayed a reduced localness of interactions.

To determine whether chromatin with higher mobility exhibits altered interactions, the relationship between these quantities was analysed (Fig. 3c). There is a subtle statistically-significant positive correlation between localness of interactions and mobility. The trend is clear for the lower mobility regions, but does not continue for beads with mobilities larger than about 0.05 *µ*m^−2^ (there we see a larger spread of localness values, but this constitutes fewer than 9% of beads). This is consistent with protein binding sites having low localness values, being in higher density regions, and having lower mobility. Note that this is an “active” locus (even in *Pax6* OFF cells, several of the surrounding genes are active) so different chromatin environments at nearby repressed or inactive regions (which are not represented in the present model) might also affect interaction locality.

### Locus conformations change on a timescale of minutes

Recent single cell microscopy^4^ and Hi-C data^6^ have revealed a striking variability in chromosome structure (in terms of chromosome domains) between cells within a population. Consistent with this, our previous work on *Pax6* shows a wide range of locus conformations across a simulated population, particularly for the *Pax6* HIGH cell line. A key question for understanding how this variation plays a role in gene regulation is whether a single live cell viewed over time would display this wide variation, or if instead each cell adopts a structure which stays relatively static once established (i.e., on long time scales similar to a cell cycle).

For each simulation the full details of the locus conformation can be accessed providing 3D coordinates of all *N* beads, and therefore it is possible to track how each moves in time. To explore this in a pragmatic manner, which is closer to what might be realised experimentally, another approach is to track a smaller number of points within the locus. Here we considered the three FISH probe regions located at the *Pax6* gene and upstream and downstream regulatory region, (URR and DRR) and taking the three pairwise separations of these probes as a description of the locus configuration. These separations are represented by a point in a 3-dimensional space (Fig. 4a). In this way, we can write a 3-dimensional vector **X** = (*x*_UP_, *x*_UD_, *x*_PD_), where the components *x*_UP_, *x*_UD_, and *x*_PD_ are the separations of the URR and *Pax6* probes, the URR and DRR probes, and the *Pax6* and DRR probes respectively. In Fig. 4b each point represents a single instant in time in one of our 20 repeat simulations of the locus in *Pax6* ON cells. The trajectory of **X** as it moves through this “configuration space” can be overlaid on the scatter plot of all data points (Fig. 4c; see Supplementary Fig. S6 for examples from *Pax6* HIGH and OFF cells).

**Fig. 4.**
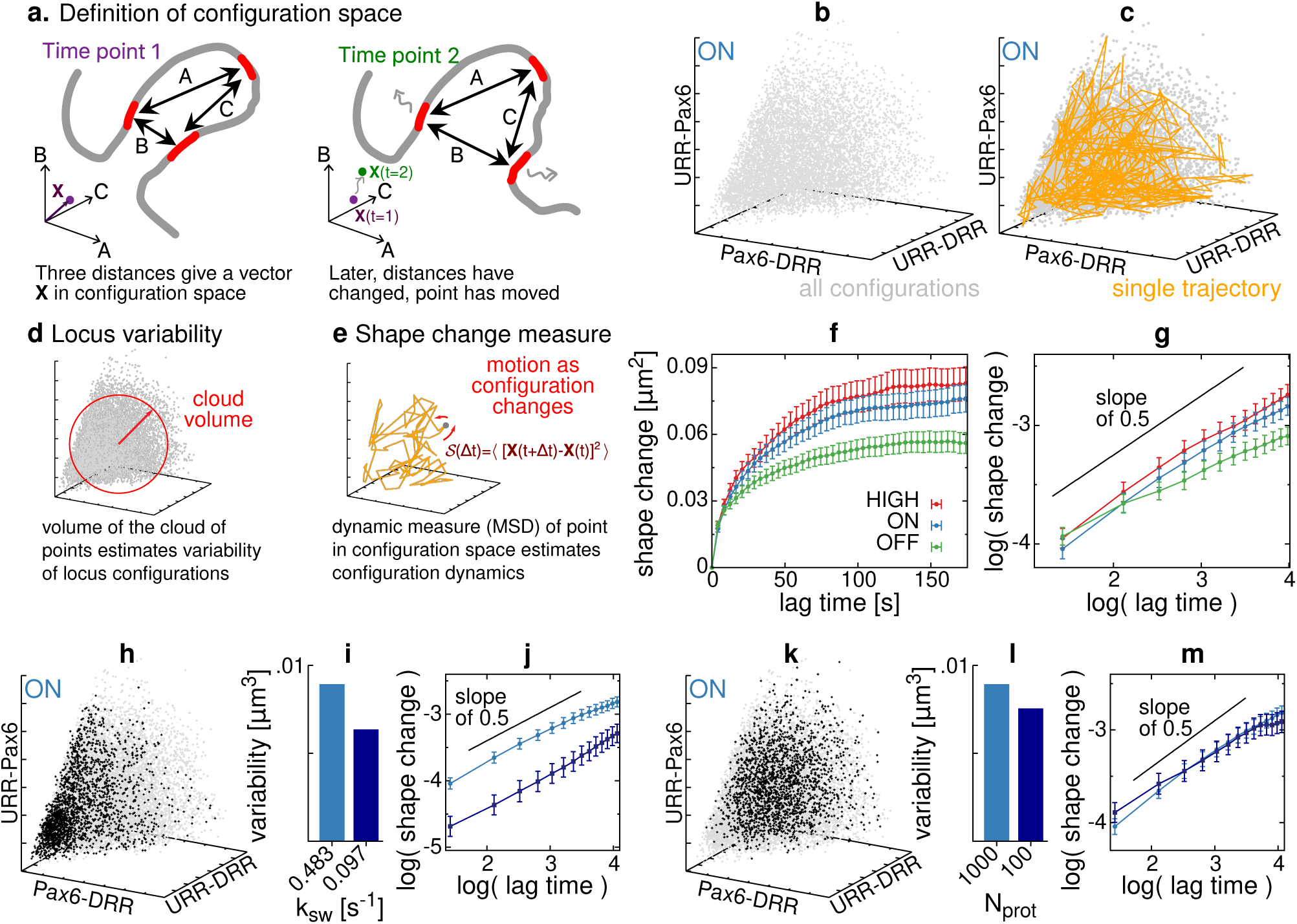
Dynamics of locus conformation. **a**. Locus configuration can be characterised quantitatively using the three distances between three regions covered by FISH probes (*Pax6*, URR and DRR). These three distances can be thought of as a point a 3-dimensional configuration space. Then as the locus configuration changes in time, this point moves in configuration space. **b**. Scatter plot showing the configurations adopted by the locus in *Pax6* ON cells. Each point represents a single time point; data are shown for all 20 repeat simulations. **c**. Same scatter plot as panel a, but overlaid (coloured line) is a trajectory from a single representative simulation; i.e., the path through configuration space is shown. The time interval between points along the path is equivalent to about 4 s, and the total duration of the simulation is about 27 minutes. **d**. The size of the cloud of points in configuration space represents the variability of the locus. **e**. The trajectory of a point through configuration space can be used to quantify how quickly the locus conformation changes via a ‘shape change parameter’. **f**. The shape-change parameter is plotted as a function of lag time for each cell line. Error bars show the standard error in the mean. **g**. The same data are shown on a logarithmic scale. The black line has a slope of 0.5 indicating sub-diffusive motion in configuration space. **h**. A scatter plot of configurations is shown for the *Pax6* ON cells from simulations but with a reduced protein switching rate (black points show results for *k*_sw_ equivalent to 0.097 s^−1^, compared to grey points where *k*_sw_ ≈ 0.49 s^−1^ as in panel a). **i**. Bar plot showing how locus variability changes with different switching rates. **j**. The shape-change parameter is shown for *Pax6* ON cells from simulations with different switching rates. **k**. Similar scatter plot as panel e, but showing simulations with 100 proteins (black) and 1000 proteins as in panel a (black). Here the switching rate is *k*_sw_ ≈ 0.49 s^−1^ in both cases. **l**. Bar plot showing how the number of proteins affects locus variability (*k*_sw_ ≈ 0.49 s^−1^). **m**. The shape-change parameter is shown for *Pax6* ON cells from simulations with different numbers of proteins.

The volume of the cloud of data points shown in configuration space represents the range of different structures which the locus can adopt in the *Pax6* ON cells (Fig. 4d) and provides a metric for “locus variability”. Examination of the configuration trajectories (Fig. 4c) suggests this volume can be explored within a single simulation of duration 7.96 × 10^5^*τ*, equivalent to roughly 27 minutes which is considerably shorter than a typical cell cycle duration. This indicates that much of the structural variability of a locus can be exhibited within a single cell.

To examine this more quantitatively a shape-change parameter *S*(*t*) was defined which tracks the mean change in locus configuration over a lag time *t*. Mathematically, this is the mean squared displacement of **X** as it moves through configuration space (Fig. 4e, see Supplementary Information for further details). As expected, *S*(*t*) saturates at large times which means that the locus explores a finite volume within configuration space (Fig. 4f). That saturation occurs within about 100-200 s implies that the locus explores all of its configuration within this time. The short time behaviour, where *S ∼ t*^*α*^, gives information about the dynamics of the locus structure: all three cell lines show *α* < 1, indicating sub-diffusive behaviour in configuration space (Fig. 4g). *Pax6* ON and HIGH cells have *α ≈* 0.5, with HIGH cells in general having larger *S* indicating that the configuration changes more quickly. Sub-diffusion is expected due to the polymeric nature of the chromosome; chromatin loops which are transiently stabilised by protein bridges or loop extruders might also be expected to act as dynamical traps, further inhibiting the motion. The *Pax6* OFF *S*(*t*) curve shows a smaller exponent *α*, perhaps reflecting the very different pattern of protein binding sites and CTCF in those cells.

How well do simulations reproduce the dynamic behaviour of real locus structures *in vivo*? It is likely that the dynamics depend heavily on the model parameters; for example, the number of proteins and the rate at which they switch between binding and non-binding states, or the number of extruders, their binding rate and extrusion speed. So far, we have used parameters within biologically reasonable ranges that have been optimised to best predict static (fixed) single cell and population level measurements^7^. However, it is possible that there are distinct sets of parameters which give similar static predictions, but with different dynamic behaviour (for example, in Ref. 34 it was shown that for the loop extruder model the ratio of extrusion speed and unbinding rate has a larger effect on predicted domains than each parameter alone). Although direct experimental measurement of the parameters used in the model remains challenging, we can use the simulations to examine how varying the different parameters affects the simulations.

First, simulations with a different protein switching rate were performed for the *Pax6* ON cell line (Fig. 4h). Specifically, the original switching rate *k*_sw_ = 10^−3^ *τ*^−1^ ≈ 0.48 s^−1^ (grey points) was compared to a reduced switching rate of *k*_sw_ = 2 × 10^−4^ *τ*^−1^ ≈ 0.097 s^−1^ (black points). A lower switching rate leads to a smaller volume being explored in configuration space (equivalently, the simulations show a reduced variability, Fig. 4i). Particularly, configurations where one or more of the probe pair separations is small are favoured over extended configurations (consistent with this, the mean size of the locus reduces). The shape-change parameter *S*(*t*) further reveals that the locus changes configuration more slowly (Fig. 4j). These observations can be rationalised since reducing the protein switching rate leads to an increase in the size and longevity of the protein clusters^25^. Therefore, protein stabilised chromatin loops are more likely to form for extended times, slowing the dynamics and favouring more compact configurations. Longer lived and larger clusters might also favour the formation of “rosettes” made of multiple chromatin loops where multiple binding sites on the chromatin come together, again reducing the overall size of the locus.

If the switching rate is kept constant, but instead the *number* of proteins is decreased, there is also a small decrease in variability, but to a lesser degree (Fig. 4l). The configurational scatter plot (Fig. 4k) reveals that in this case, configurations where the probe pair separations are small are disfavoured: the mean size of the locus *increases*. This is consistent with a reduced number of proteins leading to a lower likelihood of protein-stabilised loops forming. Unlike when altering the switching rate, here there is very little change in the configurational dynamics (Fig. 4m). These results further support a complex and non-trivial interplay between the different ingredients in the model. A decrease in locus variability can be accompanied by *either* an increase *or* a decrease in mean locus size and does not necessarily lead to a change in the locus configurational dynamics. We note that these changes to the number of proteins or the switching led to poorer agreement with the experimental data in fixed cells (CaptureC and FISH), but nevertheless provide insight into the biophysical mechanisms underlying the dynamics of a gene locus.

### Dynamics of enhancer-promoter collisions

A complementary approach to characterise locus configurations is to examine specific promoter-enhancer interactions, and track these in time. For example, the duration of a promoter-enhancer interaction (or ‘collision’) and the time interval between such interactions (i.e., collision interval) can be measured (Fig. 5a). Extraction of this data shows how often the *Pax6* promoter region interacts with each of the distal enhancer regions (Fig. 5b, and see Supplementary Information for details; note this is a coarser measurement than the precise interaction profiles shown in the simulated CaptureC data). For a long uniform polymer, the mean time between collisions of any two internal points only depends on their separation. All three of the main model ingredients will alter this. Loop extruders can actively bring different polymer regions together, and so the collision interval will depend on the pattern on CTCF sites (loop anchors). The collision interval for a given pair of sites will also depend on the protein mediated stabilisation of other nearby loops, so binding site positions across the whole locus play a role, and so does the heteromorphic polymer structure between the pairs of sites, which alters the chromatin flexibility locally. All of these factors differ between the cell lines. In general, collision intervals are largest in *Pax6* OFF cells, where there are few protein binding sites or H3K27ac (open) regions. The *Pax6*-URR collision interval is similar in *Pax6* ON and HIGH cells, presumably because the pattern of binding sites and H3K27ac in the intervening region is similar in the two cell lines. The *Pax6*-DRR collision interval, however, is larger in *Pax6* HIGH cells than ON cells probably because in *Pax6* HIGH cells there is a large region of disrupted chromatin (marked by H3K27ac) between these two sites.

**Fig. 5.**
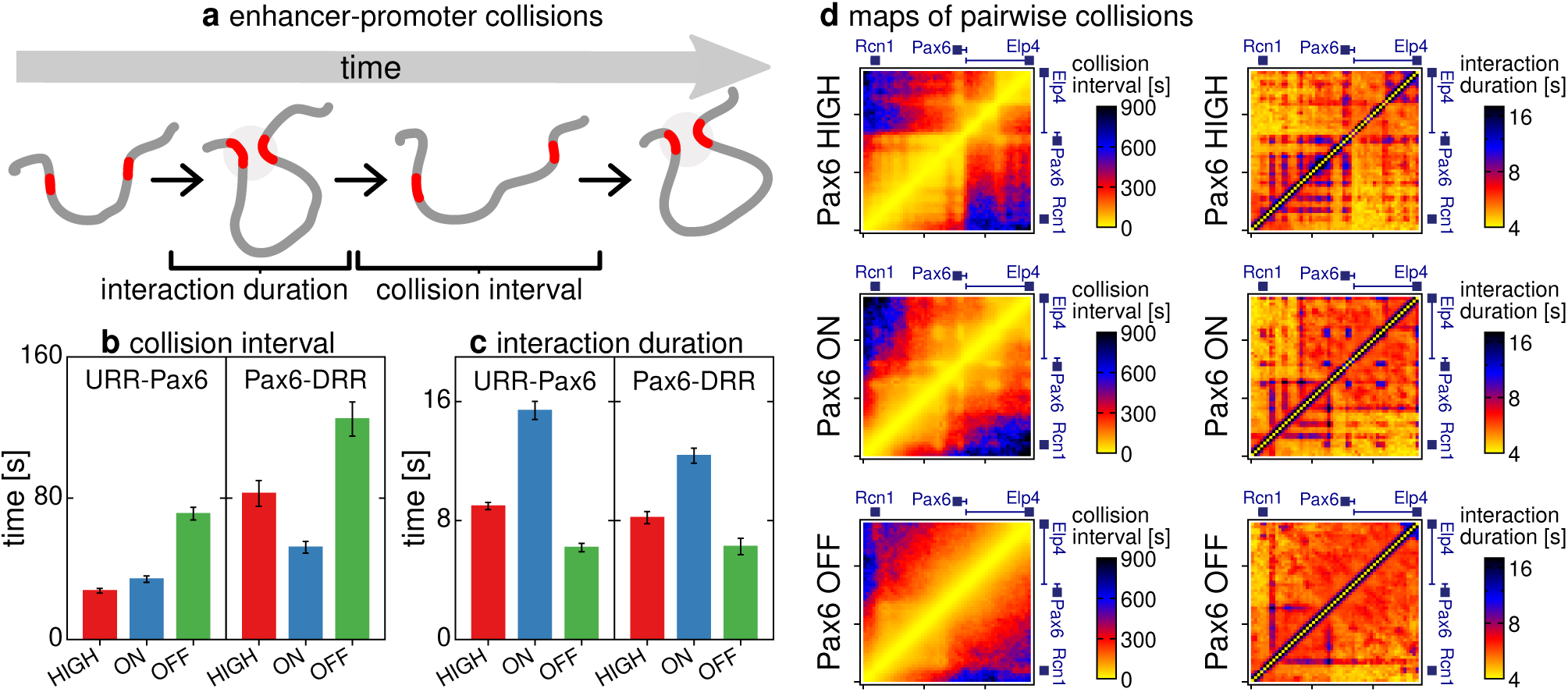
Enhancer-promoter collisions. **a**. The time interval between successive promoter-enhancer interaction events, and the duration of such events can be measured. We considered a region covering the *Pax6* promoters (chr2:105,509,500-105,520,500 mm9 reference genome) and regions covering the upstream and downstream regulatory regions (chr2:105,444,500-105,457,500 and chr2:105,629,500-105,640,500 respectively), defining a collision as when the centre of mass of the two regions has a separation of 6*σ ≈* 106 nm or less (see Supplementary Information for details). **b**. Bar plot showing the collision interval for each enhancer in the three cell lines. Error bars show standard error in the mean. **c**. Similar plots showing the mean duration of interaction (collision) events. **d**. Maps showing the mean collision interval (left) and mean interaction duration (right) for pairs of 10 kbp windows tiled across the locus.

The duration of promoter-enhancer interactions (Fig. 5c), is an order of magnitude smaller than the collision interval. The interaction duration is longest in the *Pax6* ON cells where there are more protein binding sites within the regulatory regions than in the other cell lines. The fact that the difference between, e.g., interaction durations in *Pax6* OFF and ON cells is at most about two-fold is perhaps surprising. This suggests that the stabilisation of loops by protein clusters is modest, and might be perturbed by varying, for instance, protein number and switching rate.

Collision intervals and interaction durations were also obtained from pairs of regions across the locus. Maps of these measurements (Fig. 5d) are reminiscent of Hi-C data, but instead show information on dynamics which would be inaccessible experimentally.

### Gene expression or chromatin topology perturbation only weakly affects locus conformation

Given the apparent links between locus configuration, gene activity, and dynamics, we performed new CaptureC and FISH measurements on *Pax6* HIGH cells after treatment with drugs to perturb either transcription or topology. Alpha amanitin was used to inhibit transcription through the selective degradation of elongating polymerases^35,36^. Surprisingly, but as observed previously^37^, the CaptureC profiles looked very similar to those from untreated cells, with similar interaction peaks (Fig. 6a). Although interaction peaks appeared slightly higher in the treated cells these results suggest that chromatin interactions are not driven by the process of transcriptional elongation. Similarly, an examination of locus compaction using FISH with probes located at the *Pax6* promoter and the promoters of the two neighbouring genes, *Rcn1* and *Elp4*, also showed that there was no significant change to the distributions of probe separations after treatment (Fig. 6b). Together these results suggest that inhibiting transcription, *per se*, does not affect the structure at the scale of a locus, but that interactions which occur frequently (visualised by CaptureC peaks) are enhanced.

**Fig. 6.**
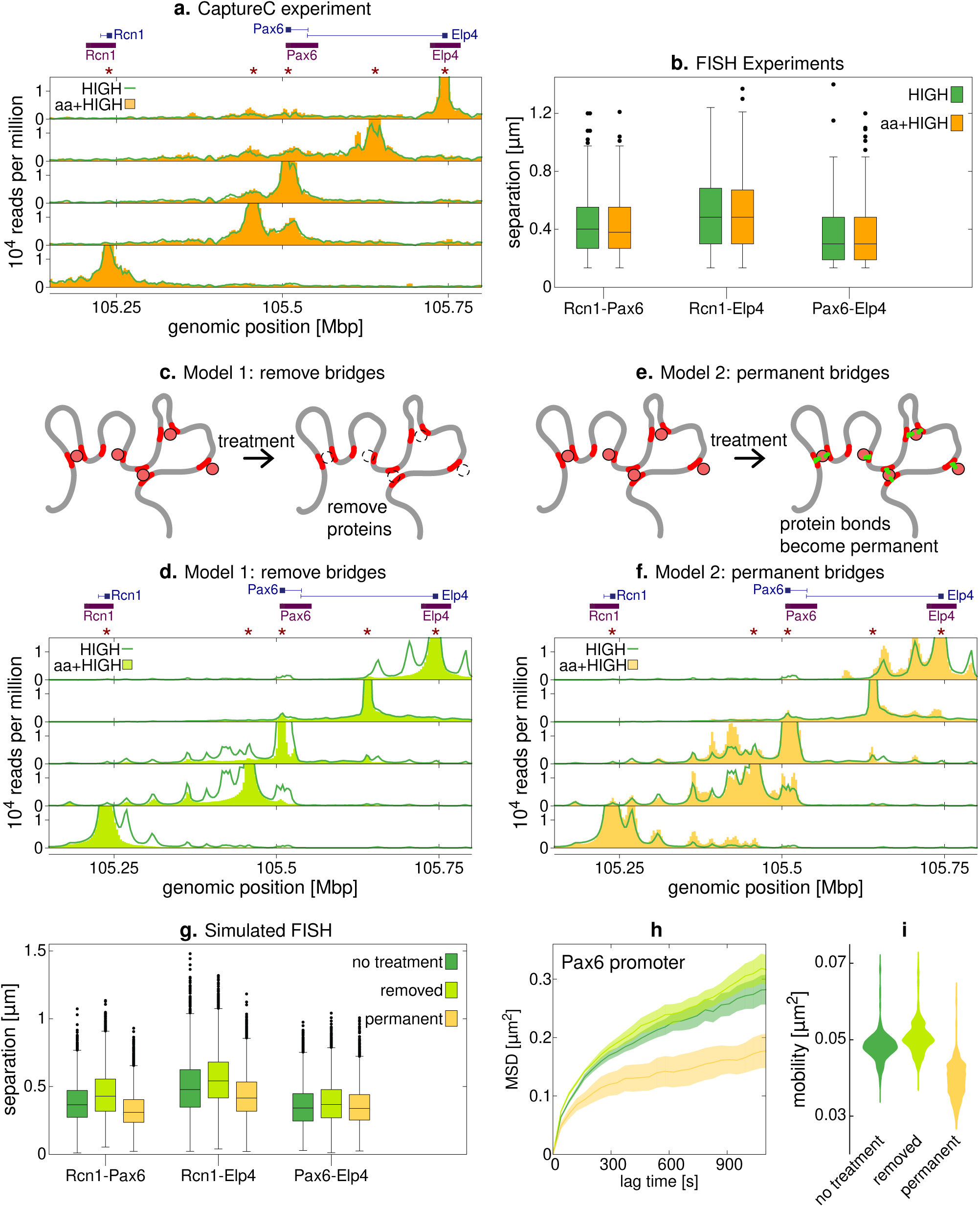
Perturbing transcription weakly affects locus conformation. **a**. CaptureC data from *Pax6* HIGH cells in an experiment with alpha amanitin treatment is shown (yellow bars) alongside similar data from untreated cells (green line). **b**. FISH data for alpha amanitin treated and untreated *Pax6* HIGH cells. Probes were positioned at *Pax6* and at the promoters of the two adjacent genes *Rcn1* and *Elp4* (purple bars in panel a and probe set 2 in Fig. 1b). Box plots show the distributions of each separation; between 129 and 146 measurements were made for each case. A Mann-Whitney U test could not reject the null hypothesis that the treated and non-treated distances were drawn from the same distribution (*p* > 0.05 for all three pairs of probes). **c**. One possible model for alpha amanitin treatment is to remove all proteins, since these could represent complexes containing RNA polymerase (see text). **d**. Simulated CaptureC results for *Pax6* HIGH cells from simulations where all protein binding sites were removed (lime green bars). Equivalent results without the simulated treatment are also shown (green line). **e**. An alternative model is where at the point of alpha amanitin treatment all protein-chromatin bonds are fixed (see text and Supplementary Information). **f**. Simulated CaptureC results for *Pax6* HIGH cells from simulations where all protein-chromatin bonds were fixed at the point of treatment (yellow bars). **g**. Plot showing simulated FISH measurements for the two different alpha amanitin treatment models, as well as the untreated case. **h**. The MSD of the *Pax6* promoter FISH probe is shown for the three cases. **i**. The mobility is calculated for each chromatin bead within the locus. The distribution of mobility values is shown as a violin plot.

To interpret these results, we considered how the effects of transcription inhibition could be modelled within the simulations. A loss of transcription could be viewed as a loss of bridge protein binding, since these model proteins represent complexes of polymerase and transcription factors (Fig. 6c). However, a simulation where all binding sites were switched off showed a dramatic loss of many of the interaction peaks in the simulated CaptureC profiles, and a general decrease in interactions across the locus (Fig. 6d) resulting in an effect *opposite* to that observed in the alpha amanitin treatment experiments.

In the HiP-HoP framework, the bridging complexes continually switch between a binding and a non-binding state^25^ to model active chemical reactions (e.g., post-translational modifications) and provide a realistic turnover rate of proteins in active foci. Another possible effect of alpha amanitin treatment is that it interrupts such reactions, through, for example, the transcription cycle of polymerase, which then remains stuck in an initiation state. A modified simulation was used to test this idea: all protein-chromatin bonds were “fixed”, so that from the point of simulated alpha amanitin treatment, any protein which is bound to one or more chromatin bead remains bound (Fig. 6e, see Supplementary Information for details). This led to an increase in interactions at many peaks in the simulated CaptureC data (Fig. 6f), a situation that is closer to what was observed experimentally than in the scenario where bridging proteins were removed.

Simulated FISH measurements showed a different trend for each of the two treatment models (Fig. 6g). Removing proteins led to an increase in probe pair separations, whereas fixing protein-chromatin bonds led to a decrease. In the simulations the dynamics of the locus also change for both of the alpha amanitin treatment scenarios, as measured via the MSD of FISH probes and mobility (Fig. 6h,i). Bridging protein removal leads to an increase in dynamics, whereas the protein fixing model leads to a decrease. Particle-tracking experiments where histone proteins were labelled fluorescently indicate that chromatin is typically more dynamic after alpha amanitin treatment^38^.

Since neither model can fully predict both FISH and CaptureC data we suggest that alpha amanitin treatment in reality results in a mixture of these effects: loss of some bridging interactions (e.g., mediated by complexes involving elongating RNA polymerase) resulting in increased mobility and local decompaction, but some other bridging interactions bringing together enhancers and promoters (e.g., mediated by transcription factors, or involving paused polymerases) may be stabilised, resulting in more pronounced CaptureC peaks. In a complementary experiment, *Pax6* HIGH cells were treated with bleomycin to perturb the local topological chromatin state. This drug nicks the DNA, releasing superhelical tension so that the DNA is torsionally relaxed^36^. After perturbation, CaptureC profiles looked very similar to those from untreated cells whilst single cell microscopy revealed no significant difference between the FISH probe separations with and without treatment. From this experiment it appears that at this scale of analysis superhelical tension, which has dramatic effects on larger scale chromatin structure^36^, does not affect chromatin architecture within the locus.

## Discussion

A strong motivation for this study was to understand the apparent contradiction of stable promoter-enhancer loops with a need for regulatory elements to continuously sample a locus for regulatory inputs throughout a cell cycle. Although the HiP-HoP model was originally developed to provide chromatin structure predictions in fixed cells, it became apparent that it could be developed to provide insight into locus dynamics, importantly in both cell populations and in a single cell.

Chromatin mobility can be determined by extracting the mean-square displacement (MSD) from the simulation. These analyses show that locus dynamics depended strongly on the size of the chromatin segment being analysed, or in other words the ‘probe’ used to track the locus. Chromatin neighbourhoods (35-40 kbp, typical of FISH probes) did not show strong differences in mobility either at different points within a locus, or in different cell lines (Figs. 2b-d) whilst small (1 kbp) probes, revealed a strong variation in their movement across the locus. This was dependent on two factors (i) local chromatin fibre structure (marked by H3K27ac) and (ii) strong bridging interactions between regulatory elements. Typically, disrupted chromatin^20^ was more dynamic than compact chromatin suggesting that regions that have a heterogeneous pattern of nucleosome binding sites that are not able to package tightly will be more dynamic. In contrast, regions that are enriched in protein binding, often marked by ATAC-seq peaks, and that form looping interactions are less dynamic. Consequently, loci that have a large number of ATAC-seq binding sites are likely to be less dynamic. However, active biding sites tend to be embedded within H3K27ac regions, leading to a balancing act between bridging-mediated slowing down and acetylation-mediated speed up of chromatin motion.

This combination of experiments and modelling suggests that the position and size of a ‘probe’ will determine the observed dynamical properties. This will be critical when designing future live-cell imaging experiments and could explain some previous, seemingly contradictory, results: in live-cell imaging where single enhancers and promoters were labelled using a CARGO-dCas9 assay where guide RNAs spanned a ∼ 2 kbp region, these became more mobile (faster dynamics) when active^39^. On the other hand, experiments using an ANCHOR/ParB DNA-labelling system to track the motion of a gene promoter found that the motion becomes constrained upon activation^40^, with faster dynamics restored if transcription initiation is inhibited. Also, several studies where multiple points across the genome were tracked (e.g., with histone proteins labelled at low concentration) revealed that chromatin dynamics tend to increase as activity is *reduced* (either by transcription inhibition, serum starvation, or when PolII is degraded^8,31,38^). These observations can be reconciled if, as suggested by our model, gene activation leads to chromatin decompaction (increasing chromatin mobility) in some regions, but additional looping or bridging (decreasing mobility) in others.

This study can begin to introduce the concept of *sampling* between regulatory elements within a locus, i.e., where the interactions between these elements continually change in time. Although there is clear movement within a locus, the concept of promoter-enhancer interactions has given rise to a notion that chromatin mobility is a limiting factor controlling locus sampling. In contrast these results indicate that for a typical locus every possible interaction event can be sampled within tens of minutes (Fig. 4). This suggests that even though the promoter-enhancer contacts revealed by 3C techniques are enriched above a background of interactions, there is continuous sampling. Unlike in laboratory-based experiments, data providing insight into how the configuration of the locus changes in time can be extracted from the simulations. Importantly, we found that within a given simulation trajectory the locus was able to explore all of its configuration space. While we expect these dynamics will depend strongly on the model parameters, our results show that a simulation using the set of parameters validated by comparison with fixed-cell structures suggests the locus can visit a large part of its configuration space within a few minutes. By defining a shape-change measure, we could also show how varying the model parameters changed these ‘configurational dynamics’. The dependence on the number of active proteins and the rate at which these switched between binding and non-binding states^25^ is subtle. Importantly, we found that different changes to the parameters which led to the same change to static properties (e.g., a decrease in locus variability) do not necessarily lead to the same change to dynamic properties. We note that the configurational dynamics will also depend strongly on the size of the region under consideration. Here we saw the ∼ 200 kbp *Pax6* region varying widely within a simulation time equivalent to a few minutes; we would expect, e.g., a 1 Mbp TAD to take much longer to re-organise.

We then found that treating cells with alpha amanitin to inhibit transcription did not greatly alter the locus structure, only leading to a small increase in the height of interaction peaks in CaptureC data. A naive model where transcription inhibition was mimicked by abrogating protein binding did not reproduce these observations. The CaptureC results are more consistent with a model where chromatin-protein-bridges are instead fixed in place. Our experiments stand in contrast with recent Hi-C, HiChIP and OCEAN-C data showing that acute degradation of PolII led to a small *decrease* in local looping interactions^41^; in another study, using MicroC in mouse ES cell, it was found that transcription inhibition using triptolide or flavopiridol did not significantly affect promoter-enhancer interactions, but the intensity of other features of the interaction maps (stripes) were reduced^42^. Older work showed that alpha amanitin treatment did not disrupt enhancer interaction hubs^37^, and that elongation inhibition via treatment with DRD or initiation inhibition by heat shock did not disrupt foci of PolII, but the latter caused active promoters to dissociate from foci^43^. Together this points to a complicated relationship between transcription and chromatin contacts, and to subtleties in the action of inhibitors like alpha amanitin which are not fully understood; nevertheless, our simulation scheme enabled us to make predictions about the effect on the dynamics that is expected in different scenarios.

These simulations of the *Pax6* locus give insight into the factors which affect chromatin dynamics and suggest that for a locus of this size, a single cell would show the same level of variability in time as is observed from cell-to-cell across a population. It would also be interesting to study dynamics in simulations of larger chromosome regions or with additional model ingredients, which might be able to shed light on the origins of some of the other experimental observations made in live cells, such as of correlated chromatin motion^8,31^, of dependence on nuclear structural proteins like lamins^44^, and of gel-like features of chromatin^45–47^. Further models could extend the HiP-HoP framework to include, for example, repressive proteins which compact DNA, interactions with the nuclear lamina, chromatin self-interactions, or take into account proteins such as SAF-A which are thought to form an RNA-dependent gel constraining chromatin motion^45^.

## Methods

### HiP-HoP simulations

The HiP-HoP model was used as previously described^7^. In brief, a chain of beads connected by springs represented a chromosome region, with 1 kbp of DNA per bead. Diffusing spheres represented complexes of proteins which bind at active chromatin sites, and these switched back and forward between a binding and a non-binding state with rate *k*_sw_ = 10^−3^ *τ*^−1^ (unless otherwise stated), where *τ* is the simulation time unit. DNA accessibility data (ATAC-seq) was used to identify binding sites. Loop extrusion was modelled by introducing additional springs between adjacent beads in the chain, which were then moved at regular time intervals to extrude a loop. Extruders were initiated at random positions, moved at a rate *k*_ex_ =2 bp *τ*^−1^, and were removed with rate *k*_off_ = 2.5 × 10^−5^ *τ*^−1^. ChIP-seq data for CTCF and Rad21 binding were used to identify loop anchor sites, and extrusion was halted at loop anchor sites in a direction dependent manner. In some regions of the bead chain, additional springs were added to “crumple” it into a more compact structure. Histone modification data (H3K27ac ChIP-seq) were used to identify regions which do not have the crumpling springs. The dynamics were evolved via a Langevin dynamics scheme (implicit solvent) using the LAMMPS molecular dynamics software^48^. As detailed in the text we simulated a 40 Mbp (40,000 bead) fragment with 10 copies of a 3 Mbp region around *Pax6* (chr2:104,000,000-107,000,000 mm9 reference genome) placed along it. Periodic boundary conditions were used, and the system size was chosen to give a roughly realistic chromatin density. We used data from three different mouse cell lines, and a mixture of loci from each was included in a given simulation. Each of 6 independent simulations were run for 1.5 10^6^ simulation time units, with the first 7 × 10^5^ *τ* discarded to allow the system to each a steady state where the polymer configuration was locally at equilibrium. We generated 20 trajectories (representing single cells) for each cell line, extracting a total of 8000 configurations per cell line. Full details of the simulations, interaction potentials, and all parameters are given in Supplementary Information. The input data were previously published^7^ and are available at GEO: GSE119660, GSE119656, GSE119659, GSE119658, GSE120665, and GSE120666. Full details of the data analysis are given in Supplementary Information.

### Mapping simulation length and time scales to real units

Simulation length and time units were mapped to physical units by comparing with experimental data. To estimate the length unit, for a given pair of FISH probes a distribution of separations was obtained from simulations and experiments^7^, and compared using the Kolmogorov-Smirnov statistic (the smaller this statistic the closer the two distributions). We obtained nine such distributions (three distances between *Pax6*, URR and DRR probes in each of three cell lines), and used the mapping which minimised the average of the nine Kolmogorov-Smirnov statistics. This gave an estimate for the length unit of *σ ≈* 17.6 nm. To map the simulation time unit, we calculated an MSD for every chromatin bead in every simulation, and obtained an average. This was compared with data from motion tracking experiments in Ref. 29 where MSDs were obtained for several chromosome regions in *Saccharomyces Cerevisiae*. We used a linear fit to find the mapping which minimised the difference between the simulated and experimental MSD curves. This led to an estimate for the simulation time unit of *τ* ≈ 2.07 × 10^−3^ s.

### Alpha amanitin and bleomycin treatment experiments

*Pax6*-HIGH cells (also known as *β*-TC3 cells) were isolated from a mouse insulinoma^49^ and were cultured in Dulbecco’s Modified Eagle Medium (DMEM) (ThermoFisher) supplemented with 10% fetal calf serum and 1% Penicillin-Streptomycin at 37°C in 5% CO_2_. No cell authentication was performed, and the sex of the cell line is not known. Transcription was blocked by adding alpha amanitin (50 *µ*g ml^−1^) to cells. For chromatin nicking, cells in six-well plates (for CaptureC) or on slides (for FISH analysis) were treated with bleomycin (100 *µ*M) for 10 min.

### CaptureC experiments

For *Pax6*-HIGH cells which had been treated with alpha amanitin or bleomycin, NG CaptureC was performed as previously described^27,28^, but with the following alterations. Two replicates of 5 × 10^6^ cells were processed for each case, fixed with 2% formaldehyde, and lysed with standard 3C lysis buffer for 15 min before snap freezing. Cells were further lysed by resuspension in water, and then in 0.5% SDS for 10 min at 62°C. Each replicate was split between three tubes, re-suspended in 800 *µ*L 1 × *DpnII* buffer (NEB) with 1.6% Triton X-100, and digested with 3 sequential additions of 750 units *DpnII* enzyme at 37°C with 1200 rpm shaking over 24 hours. Samples were heat inactivated at 65°C for 20 min, and 3 samples from each replicate combined into 7 mL with 1 × T4 DNA Ligase Buffer (NEB), with 1% Triton X-100, and 12,000 units of T4 DNA ligase at 16°C overnight. Samples were treated with Proteinase K overnight at 65°C and RNase A/T1 (ThermoFisher) for 1 hour at 37°C, before a standard Phenol/Chloroform extraction and ethanol precipitation was performed. Complete digestion and ligation was assessed by gel electrophoresis.

Purified 3C DNA from each sample was sonicated to 200-400 bp with a Soniprep 150 probe sonicator at 4°C and purified with a standard Ampure XP Bead protocol (Beckman Coulter) using a 1/1.5 DNA to bead ratio. Two Illumina sequencing libraries were prepared per capture pool replicate, with 6 *µ*g of starting DNA in each, and generated using NEBNext DNA Library Prep Kit (NEB). Samples were indexed with unique barcodes using NEBNext Multiplex Oligos for Illumina (NEB). Two separate capture pools were designed to the following *Pax6* locus elements as in Ref. 7 (a list of targeted restriction enzyme fragments is given in Supplementary Table 1). Capture oligos were designed to each end of the targeted *DpnII* fragments^28^, and each synthesised as a separate 4 nM synthesis, with a 5′ biotin label on a 120 bp Ultramer (IDT). Capture oligos from each of the two pools were mixed at equimolar amounts and pooled to a final concentration of 13 pmol in a volume of 4.5 *µ*L per sequence capture. Libraries were sized and quality controlled on a D1000 Tapestation tape (Agilent).

NG CaptureC sequence capture was performed using SeqCap EZ HE-Oligo Kit A or B (dependent on the multiplex barcode) and SeqCap EZ Accessory Kit (Nimblegen)^28^, using each of the two capture pools, with 1.5-2 *µ*g 3C library DNA per hybridization reaction. Each hybridization reaction was performed on a thermocycler at 47°C and incubated for between 66 and 72 hours. Each hybridization reaction was then bound to streptavidin beads from SeqCap EZ Pure Capture Bead Kit and washed with SeqCap EZ Hybridization and Wash Kit (Nimblegen), following the manufacture’s protocol. Hybridization reactions were split into two and libraries re-amplified using Post LM-PCR oligos (Nimblegen) and Q5 High-Fidelity DNA polymerase (NEB) directly from the beads, and then the DNA was purified using Ampure XP Bead 1/1.8 DNA to bead ratio. A second hybridisation reaction was performed as above on the re-amplified 3C libraries with two reactions pooled together (∼ 1 *µ*g in each) and incubated for 22-24 hours. Washed and re-amplified double captured libraries were sized and quality controlled on a D1000 Tapestation tape (Agilent), and paired-end sequenced on an Illumina Hi-seq 2500 or Hi-seq 4000. CaptureC data was analysed using the capC-MAP software^50^ (see Supplementary Information for further details).

### Three-dimensional DNA fluorescence in situ hybridization

Cells were grown overnight on glass slides. Slides were rinsed with PBS and fixed in 4% paraformaldehyde for 10 min. Slides were rinsed with PBS and cells were permeabilised for 10 min on ice with PBS supplemented with 0.2% triton. After rinsing, slides were stored in 70% ethanol at 4°C.

For processing, slides were dehydrated through an ethanol series and incubated with 2 × SSC supplemented with 100 *µ*g ml^−1^ RNase A (Invitrogen) at 37°C for 60 min. Slides were then rinsed briefly with 2 × SSC, dehydrated through an ethanol series and air dried. Slides were warmed by incubation in a 70°C oven for 5 min before denaturation for 1 min in 70% formamide in 2 × SSC, pH 7.5, at 70°C. Slides were then transferred to 70% ethanol on ice, dehydrated through an ethanol series and air dried before overnight hybridization at 37°C with pairs of fosmid probes (listed in Supplementary Table 2). Probes (BacPac resources) were labelled in green-500-dUTP (ENZO life sciences), digoxigenin-11-UTP (Roche) or biotin-16-dUTP (Roche). 150 ng of each labelled probe was hybridised with 5 *µ*g salmon sperm and 10 *µ*g human Cot1 DNA. Slides were washed four times for 3 min in 2 × SSC at 45°C and four times for 3 min in 0.1 × SSC at 60°C before being transferred to 4 × SSC with 0.1% Tween 20 at room temperature. Digoxigenin-labelled probes were detected using one layer of rhodamine-conjugated sheep anti-digoxigenin and a second layer of Texas red–conjugated anti-sheep (Vector Laboratories). Biotin-labelled probes were detected using one layer of FITC-conjugated streptavidin followed by a layer of biotin-conjugated anti-avidin and a second layer of FITC-conjugated streptavidin (Vector Laboratories). Slides were counter-stained with 0.5 *µ*g ml^−1^ DAPI.

### Image capture and analysis

Three-colour 3D DNA FISH slides were imaged using a Hamamatsu Orca AG CCD camera (Hamamatsu Photonics) Zeiss Axioplan II fluorescence microscope with PlanNeofluar objectives, a 100-W Hg source (Carl Zeiss) and Chroma 83000 triple band-pass filter with single excitation filters installed in motorised filter wheels (Prior Scientific Instruments). Image capture and analysis were done using in-house scripts written for Iola Spectrum (Scanalytics). For FISH, images were collected from at least 50 randomly selected nuclei for each experiment and then analysed using custom Iola scripts that calculate the distance between two probe signals.

## Acknowledgements

DM acknowledges support from European Research Council (CoG 648050, THREEDCELLPHYSICS). NG was supported by the UK Medical Research Council (MC_UU_00007/13). We also acknowledge Michael Chiang and Cleis Battaglia for useful discussions.

## Author contributions

GF performed the simulations, and GF and CAB analysed the data. AB performed the experiments, and AB and SB analysed the microscopy data. DM, NG and CAB designed the research, and all authors contributed to writing the manuscript.

