## Supplementary Material for "Transcription modulates chromatin dynamics and locus configuration sampling"

---

#### Supplementary Figures

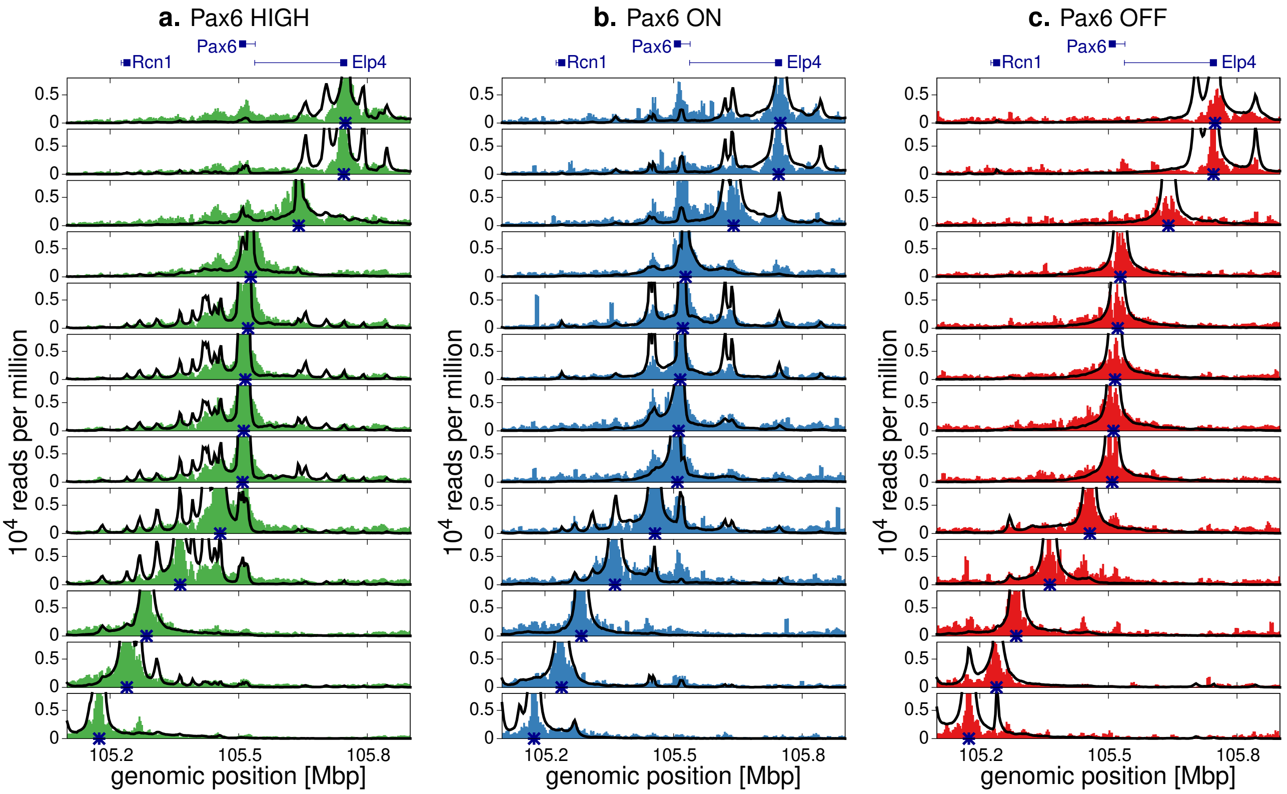


**Supplementary Fig. 1 | Simulations reproduce CaptureC data from viewpoints across the locus.** Experimental data[1] (shaded regions) are shown on the same axes as simulation results. A cross on the horizontal axis marks the position of the viewpoint. Positions of the genes are indicated above the plots. Results are shown for: **a**, *Pax6* HIGH cells; **b**, *Pax6* ON cells; and **c** *Pax6* OFF cells. A similar level of agreement between simulations was observed as in Ref. 1, where single copies of the locus were simulated in isolation.

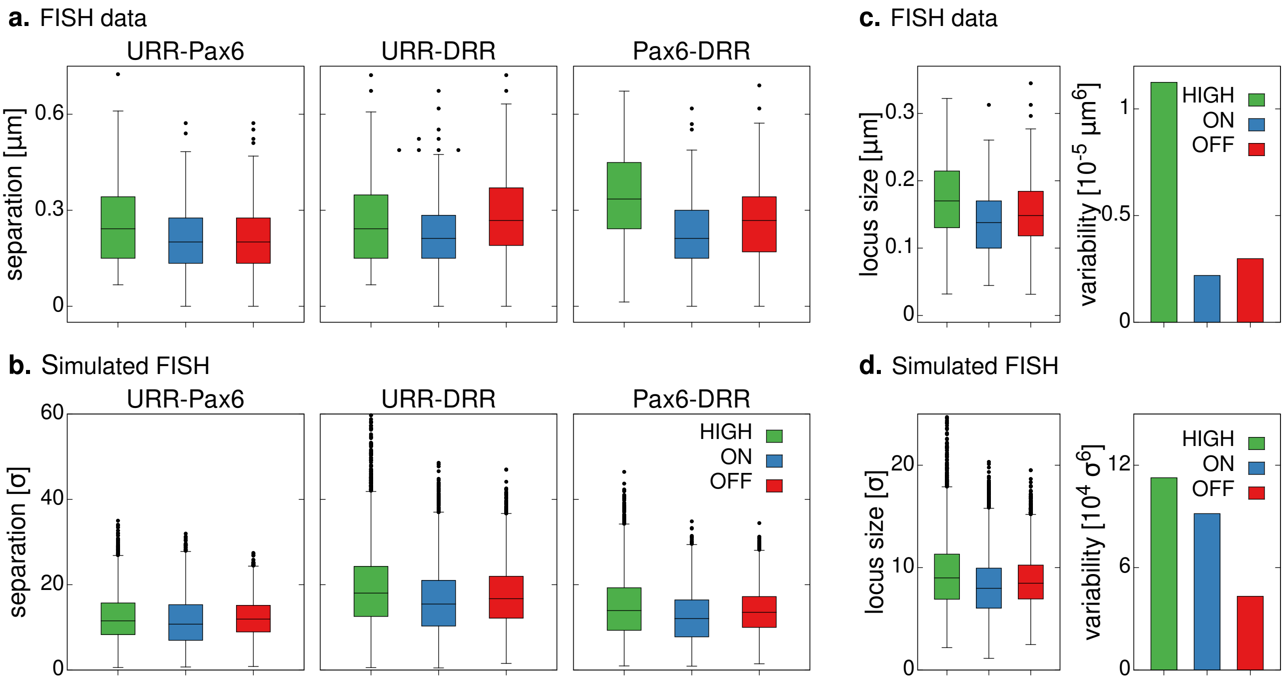


**Supplementary Fig. 2 | Simulations reproduce trends observed via microscopy. a,** FISH microscopy data was obtained from Ref. 1. Box plots show separation measurements from three pairs of FISH probes, positioned at *Pax6*, the upstream regulatory region (URR), and downstream regulatory region (URR). **b,** Equivalent measurements were obtained from simulations. Similar trends were observed as in the experimental data. Specifically, *Pax6* HIGH cells were the least compact (largest separations on average), while *Pax6* ON cells were the most compact (smallest separations on average). A similar level of agreement was observed as in Ref. 1, where single copies of the locus were simulated in isolation. Simulation length units ($\sigma$) are used; these can be mapped to real lengths by comparison with the data in panel a, as detailed in Supplementary Information. **c,** Three colour FISH microscopy allowed separations between all three probes to be measured in a given cell. This enabled the overall locus size to be determined (as defined in Supplementary Information). A measure of the cell-to-cell variability was also defined (see Supplementary Information). **d,** Similar measures from the simulations show that the trends for locus size are captured. The trend for the variability is not correctly captured in the simulations, and this is a less good agreement than in Ref. 1. This suggests that the variability is also affected by the broader surrounding environment of the locus, which is different here than in our previous work (i.e., each simulation includes multiple copies of the locus).

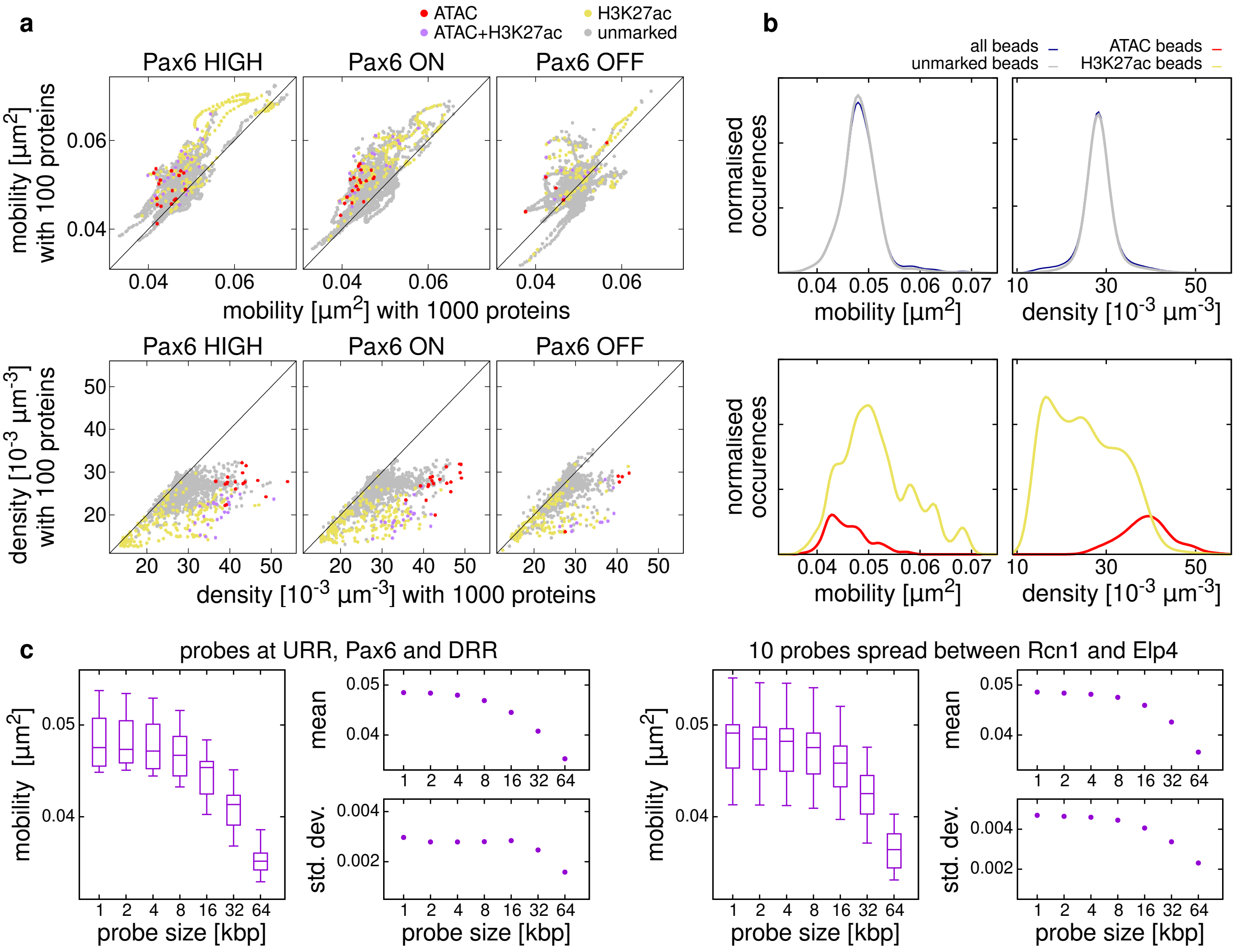


**Supplementary Fig. 3 | Factors affecting mobility a,** Model parameters affect the mobility and density at 1 kbp beads. Scatter plots compare quantities measured from simulations with 1000 proteins (as in Fig. 2) and from simulations with 100 proteins. Each point represents a chromatin bead within the locus, with colour indicating the bead properties as indicated. Top row plots compare mobilities in the two cases; points appearing above the diagonal indicate a higher mobility in the case with fewer proteins. Bottom row plots compare the density around each bead; points appearing below the diagonal indicate a lower density in the case with fewer proteins. Values are obtained from averages over 20 locus copies with 400 configurations from each for the 1000 protein case, and from averages over 10 locus copies with 200 configurations from each for the 100 proteins case. **b,** Local chromatin context affects mobility and density at 1 kbp beads. Histograms show the mobility and density for beads across all cell lines (1000 protein simulations as in Fig. 2). Values are split according to the properties of the bead (ATAC sites, H3K27ac marked, unmarked). Top plots reveal that unmarked beads show the same mobility distribution as all beads. Bottom plots show that while ATAC beads generally have lower mobility and H3K27ac beads generally have higher mobility, both types of bead can take a broad distribution of values. Density distributions show similar but opposite trends. Note that just over half of ATAC beads are also marked with H3K27ac, so it is expected that these distributions overlap. The curves are kernel density plots obtained by summing a Gaussian function for each bead; for the mobility the Gaussians have width 0.001 $\mu$m$^2$, while for the density they have width $2 \times 10^{-3} \mu$m$-3$. Curves are scaled so that the area under each is equal to the number of beads of the specified type. **c,** Probe size affects the measured mobility. We define the mobility of a probe as the MSD of the centre of mass of the beads covered by the probe at a fixed lag time of $\Delta t = 10^4 \tau \approx 20.7$ s. In the left-hand plots we consider probes of different sizes at the URR, *Pax6*, and the DRR (centred on the centre of the FISH probes used in fixed cell experiments, Supplementary Table 2). In the right-hand plots we consider 10 probes of different sizes positioned equally spaced across the locus between *Rcn1* and *Elp4*. Box plots show distributions of mobilities of all probes of a given size across all three cell lines (the same simulations as in Fig. 2). The mean and standard deviation of the distributions are also shown; both decrease as the size of the probe increases.

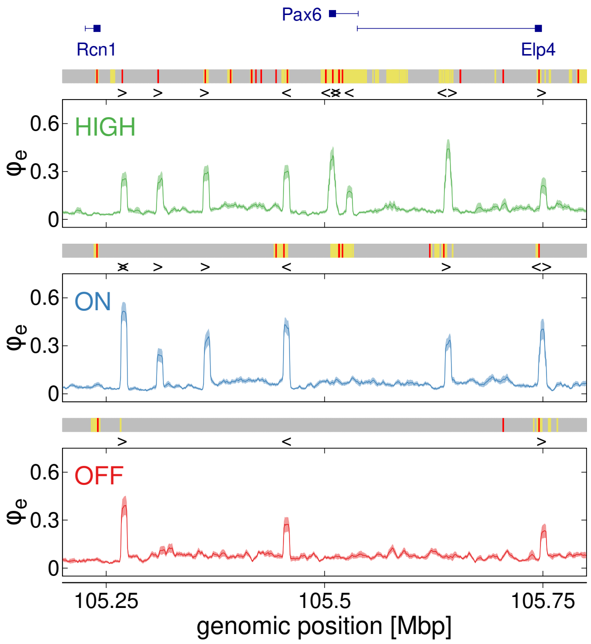


**Supplementary Fig. 4 | Extrusion probability.** Plot showing the probability that a loop extruder is positioned at a chromatin bead at any point during the simulation. The dark lines show the average over 20 copies of the locus, with the shaded region (which is often covered by the line) showing the standard error in the mean. The coloured block above each profile shows positions of ATAC (red) and H3K27ac (yellow) beads, and the chevrons indicate the positions and orientation of CTCF sites. Recall that in each copy of the locus the CTCF sites are occupied with probability determined by the peak height in the CTCF ChIP data (see Supplementary Information).

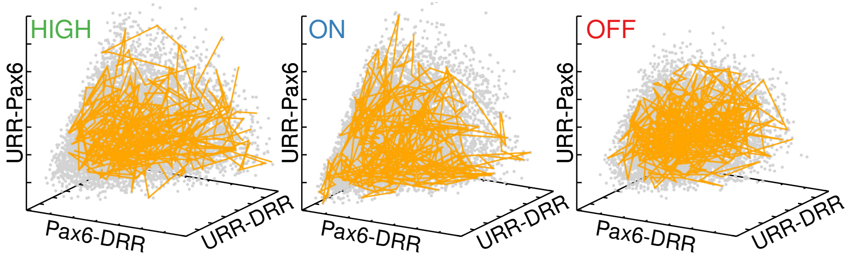


**Supplementary Fig. 5 | The *Pax6* locus explores a 'configuration space'. a,** Scatter plots showing the separations of the *Pax6* and enhancer probes in each of the different cell lines, similar to Fig. 4c. Each grey point represents a single time point; data are shown for all 20 repeat simulations of the locus in each case. The size of the cloud of points in this configuration space represents the variability of the locus. In each plot the yellow line shows a trajectory from a single representative simulation, i.e., the path through configuration space is shown. The time interval between points along the path is equivalent to about 4 s, and the total duration of each simulation is about 27 minutes.

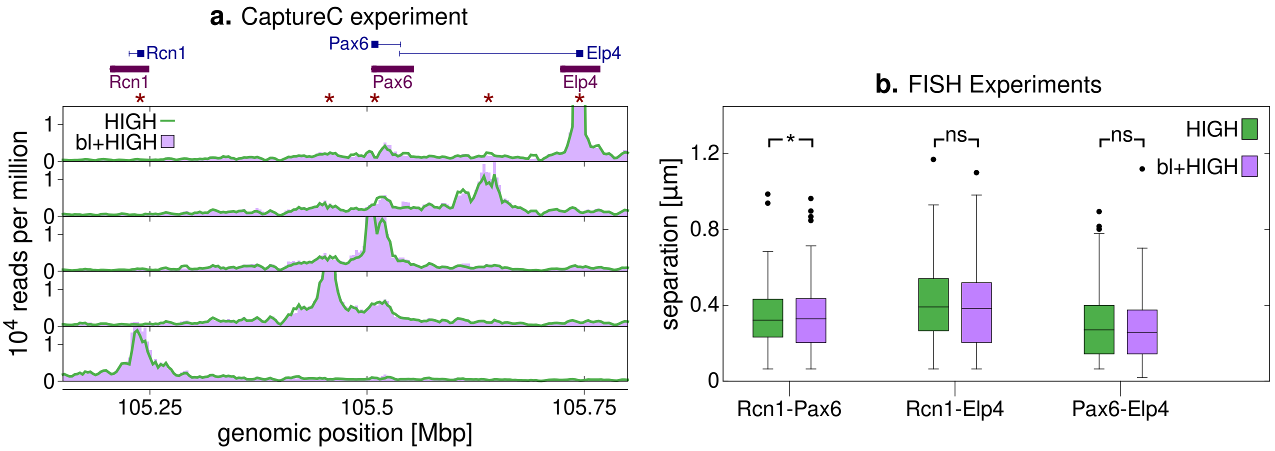


**Supplementary Fig. 6 | Bleomycin treatment relieves superhelical tension. a,** Plots showing CaptureC data from *Pax6* HIGH cells with and without bleomycin treatment. Data from five viewpoints are shown, with viewpoint positions indicated by stars above the plots. Positions of genes are also indicated. **b,** Single cell microscopy was used to measure separations of FISH probes positioned at *Pax6* and the two neighbouring genes *Rcn1* and *Elp4* (positions are indicated under the genes in panel a). Box plots show the distributions of separations for $n = 106$ measurements for bleomycin treated cells and $n = 104$ measurements for untreated cells. The * indicates that $p < 0.05$ obtained from a Mann-Whitney U test with the null hypothesis that measurements are obtained from the same distribution; 'ns' indicates there is no significant difference ($p > 0.05$).

### Supplementary Tables

| Target name | pool | restriction fragment |
|---|---|---|
| Pax6_P1 | 1 | chr2:105515339-105515903 |
| Pax6_P0 | 1 | chr2:105508737-105509119 |
| Pax6_Palpha | 1 | chr2:105521487-105522094 |
| CTCFp11 | 1 | chr2:105511309-105512303 |
| 7CE12_CTCF | 1 | chr2:105527664-105528138 |
| Elp4_pro | 1 | chr2:105744078-105745820 |
| CTCF5 | 2 | chr2:105639621-105640392 |
| CTCF6 | 2 | chr2:105456258-105457375 |
| CTCF4 | 2 | chr2:105748530-105748848 |
| CTCF7 | 2 | chr2:105363080-105364092 |
| CTCF10B | 2 | chr2:105173505-105175699 |
| Rcn1_pro | 2 | chr2:105238728-105239162 |
| e200_Enh | 2 | chr2:105284625-105285412 |

**Supplementary Table 1 |** Table showing the *DpnII* restriction enzyme fragments (mm9 coordinates) targeted in CaptureC experiments. In experiments oligos were grouped into two pools, with each pool being used in separate samples.

| Group | Name | Fosmid ID | Genomic position | Label |
|---|---|---|---|---|
| 1 | URR | WIBR1-1230I10 | chr2:105430492-105469674 | digoxigenin-11-dUTP |
| | *Pax6* | WIBR1-0322P22 | chr2:105508853-105550057 | biotin-16-dUTP |
| | DRR | WIBR1-2859L14 | chr2:105612707-105655961 | green-500-dUTP |
| 2 | *Rcn1* | WIBR1-0075F18 | chr2:105218785-105254871 | green-500-dUTP |
| | *Pax6* | WIBR1-0322P2 | chr2:105508853-105550057 | digoxigenin-11-dUTP |
| | *Elp4* | WIBR1-1660D19 | chr2:105726249-105764673 | biotin-16-dUTP |

**Supplementary Table 2 |** Table giving details of FISH probes. Coordinates are for the mm9 genome. Supplementary Fig. 2 shows data from Ref. 1 which used probe set 1. Probe set 2 was used in new experiments where Pax6-HIGH cells were treated with alpha amanitin (Fig. 6) or bleomycine (Supplementary Fig. 6).

### Supplementary Methods

#### 1 Simulation Methods

In the HiP-HoP model[1–4], a section of a chromatin fibre is represented as a chain of beads connected by springs. We then use molecular dynamics simulation methods to evolve the configuration of this polymer based on a set of phenomenological potentials which describe how the beads interact. Each bead in the chain represents a 1 kbp region of the genome. Three additional model ingredients drive the chromatin configuration: interactions with spheres representing complexes of proteins[5–7], a heteromorphic polymer structure[8], and loop extrusion[9,10].

##### 1.1 Protein bridges and the polymer model

Interactions among polymer beads are defined by a force field consisting of four potentials. First, non-adjacent beads interact sterically via a Week-Chandler-Anderson (WCA) potential given by

$$V_{\mathrm{WCA}}(r_{i,j}) = 4k_BT\left[\left(\frac{\sigma}{r_{i,j}}\right)^{12} - \left(\frac{\sigma}{r_{i,j}}\right)^{6} + \frac{1}{4}\right]\Theta(2^{1/6}\sigma - r_{i,j}), \tag{1}$$

where $r_{i,j} = |\mathbf{r}_i - \mathbf{r}_j|$ is the separation of beads $i$ and $j$ located at positions $\mathbf{r}_i$ and $\mathbf{r}_j$ respectively, $T$ and $k_B$ are the temperature of the system and the Boltzmann constant, $\sigma$ is the bead diameter, and $\Theta(x)$ is the Heaviside function [$\Theta(x) = 1$ if $x > 0$, or $\Theta(x) = 0$ otherwise]. Second, adjacent beads are connected by finitely-extensible non-linear elastic (FENE) springs with interaction energy defined by

$$V_{\mathrm{FENE}}(r_{i,i+1}) = -\frac{K_{\mathrm{FENE}}}{2}R_0^2 \ln\left[1 - \left(\frac{r_{i,i+1}}{R_0}\right)^2\right] + V_{\mathrm{WCA}}(r_{i,i+1}),$$

with $K_{\mathrm{FENE}} = 30k_BT/\sigma^2$ being the bond strength and $R_0 = 1.6\sigma$ the maximum bond length. Third, the polymer is afforded a finite bending stiffness via a Kratky-Porod potential given by

$$V_{\mathrm{BEND}}(\phi_i) = K_{\mathrm{BEND}}\left(1 + \cos\phi_i\right),$$

where $\phi_i$ is the angle between beads $i-1, i$ and $i+1$, while $K_{\mathrm{BEND}} = 4k_BT$ leads to a persistence length comparable to that of chromatin. Finally, additional springs are used to "crumple" the polymer in some regions (i.e. to give it heteromorphic properties). A harmonic spring potential is used, described by

$$V_{\mathrm{HARM}} = K_{\mathrm{HARM}}(r - R_{\mathrm{HARM}})^2,$$

with spring constant $K_{\mathrm{HARM}} = 200\varepsilon/\sigma^2$ and equilibrium bond distance $R_{\mathrm{HARM}} = 1.1\sigma$. This potential is applied between next-nearest neighbour beads ($i$ and $i+2$) which do not present ChIP-seq peaks for acetylation of histone H3 in lysine 27 (H3K27ac) which usually identifies open chromatin regions.

Protein complexes are represented by spheres of the same size as the polymer beads. They interact sterically with each other via the WCA potential as given in Eq. (1). Proteins have a strong attractive interaction with a subset of polymer beads which represent protein binding sites on the chromatin, a weak attractive interaction with chromatin beads marked by H3K27ac, and a steric (WCA) interaction with other chromatin beads. The attractive interactions are described by a shifted and truncated Lennard-Jones potential

$$V_{\mathrm{LJ/cut}}(r_{i,j}) = [V_{\mathrm{LJ}}(r_{i,j}) - V_{\mathrm{LJ}}(r_c)]\,\Theta(r_c - r_{i,j}),$$

with

$$V_{\mathrm{LJ}}(r) = 4\varepsilon\left[\left(\frac{\sigma}{r}\right)^{12} - \left(\frac{\sigma}{r}\right)^{6}\right],$$

where $r_c = 1.8\sigma$ is the cutoff distance and $\varepsilon$ is the interaction strength. For strong interactions (i.e. with protein binding sites) we use $\varepsilon = 8k_BT$, while for weak non-specific interactions we use $\varepsilon = 2k_BT$.

During the simulations proteins switch back and forward between a binding and a non-binding state with at rate $k_{\mathrm{switch}}$[7]. This represents post-translation modifications which alter the protein-DNA binding affinity (e.g. phosphorylation). When in the non-binding state the interaction between proteins and chromatin binding sites revert to the WCA potential.

##### 1.2 Langevin Dyanmics

We use the LAMMPS molecular dynamics software to perform the simulations[11]. Proteins and polymer beads move according to the Langevin equation

$$m_i\frac{d^2\mathbf{r}_i}{dt^2} = -\nabla_i U - \gamma_i\frac{d\mathbf{r}_i}{dt} + \sqrt{2k_BT\gamma_i}\,\boldsymbol{\eta}_i(t), \tag{2}$$

where $m_i$ is the bead mass, $\mathbf{r}_i$ is its position, $U$ is the total potential energy of the system and $\gamma_i$ is the friction due to an implicit solvent. The final term provides thermal noise where components of the vector $\eta_i$ are such that

$$\langle \eta_{i,\alpha}(t) \rangle = 0 \text{ and } \langle \eta_{i,\alpha}(t)\,\eta_{j,\beta}(t') \rangle = \delta_{i,j}\delta_{\alpha,\beta}\delta(t-t'),$$

where $\delta_{i,j}$ is the Kronecker delta and $\delta(t-t')$ is the Dirac delta function.

In LAMMPS, Eq. (2) is solved using a velocity-Verlet scheme. We use a time step $\mathrm{dt} = 0.01\ \tau$, where $\tau$ is the simulation time unit (see below for mapping to real times).

#### 1.3 Loop extrusion

The HiP-HoP model includes the loop extrusion mechanism thought to be performed by the cohesin complex[9,10,12]. Cohesin rings are represented by additional spring bonds between beads; the pair of beads to which the bond is applied is then moved outwards, i.e., from $(i,i+3) \to (i-1,i+4) \to (i-2,i+5)$ etc., to extrude a loop. Extruders are added at random $i,i+3$ positions with rate $k_{\mathrm{on}}$ and removed with rate $k_{\mathrm{off}}$. Their positions are stepped stochastically at rate $k_{\mathrm{ex}}$. The spring potential is given by

$$U_{\mathrm{EXTR}}(r_{i,j}) = U_{\mathrm{WCA}}(r_{i,j}) + K_{\mathrm{EXTR}}(r_{i,j} - r_0)^2,$$

where $K_{\mathrm{EXTR}} = 40 k_B T/\sigma^2$ is the bond strength and $r_0 = 1.5\sigma$ is the equilibrium length of the spring. An extruser is halted either when it meets another extruder, or when it meets a CTCF site whose direction is opposite to the extrusion direction. If an extruder is halted on one side, the position at the other side continues to be moved, extruding the loop. With this scheme, which is similar to that in, e.g. Ref. 9, there is no feedback from the 3D dynamics onto the extruder dynamics; for this reason, the positions of the CTCF sites in each simulation are the only thing affecting the extruder dynamics.

#### 1.4 Simulation Setup

As detailed in the main text, in order to perform simulations in a realistic chromatin density, we consider a larger chromatin fragment than in Ref. 1. Specifically we simulate a 40,000 bead polymer representing a 40 Mbp chromatin fragment, in a square system of side $91\,\sigma$ (with periodic boundary conditions). This gives a chromatin density of $\sim 50 bp/\sigma^3$, which once mapped to real units is the same order of magnitude as *in vivo*. To be able to obtain a useful simulation data set, instead of simulating a 40 Mbp region around *Pax6*, we simulated a 3 Mbp region, and positioned 10 copies of this region along the 40,000 bead polymer (with a 910 kbp region of "unmarked" chromatin between each). This means that a single simulation gives data equivalent to measuring 10 single cells. The limitation of this approach is that a given copy of the locus is embedded in an unrealistic background; comparison with experimental data (CaptureC and FISH, Supplementary Figs. 1,2) confirmed that this does not greatly affect predictions for interactions within the locus or locus configurations. To ensure that each copy of the locus was embedded in a similar background, in a given simulation we included a mixture of *Pax6* loci from the different cell lines, and in repeat simulations we randomly shuffled the positions of the loci from different cell lines. Results are then obtained by combining measurements for all copies of the locus in the same cell line. The majority of results presented are obtained from six independent simulations, giving in total 20 copies of the locus in each cell line. For results in Fig. 4h-m and Supplementary Fig. 3a, where we used different parameters, three independent simulations were performed (10 copies of the locus in each cell line).

As an initial condition, we arranged the 40,000 bead chains in a mitotic-like structure similar to those in Ref. 13. This was first relaxed in the absence of protein interactions but with loop extrusion. This allowed the chain to move away from its initial structure, but maintained a fractal globule-like probability of interaction as a function of genomic separation[14]. A further relaxation simulation was then done with the additional springs which "crumple" the polymer as detailed above.

In the majority of cases, each independent simulation was then run for a total time of $1.5 \times 10^6\,\tau$; we discarded the first $7 \times 10^5\,\tau$, by which time the system had reached a steady state (verified by checking that the radius of gyration of any given copy of the *Pax6* locus, or the radius of gyration of the entire polymer had stopped systematically changing, Fig. 1e). The presented results were obtained from the remaining $8 \times 10^5\,\tau$. For results in Fig. 4h-m and Supplementary Fig. 3a, where we used different parameters, shorter simulations of duration $5 \times 10^5\,\tau$ were used.

#### 1.5 Lengths, times and other units in simulations

In our simulations lengths are given in multiples of the polymer bead diameter $\sigma$; as detailed in the Methods section in the main text, we determined how this maps to a real length scale by comparison between simulated and experimental FISH measurements. (as in Ref. 1). For each pair of FISH probes in each cell line, we obtained a distribution of separations, and compared simulated and experimental distributions using the Kolmogorov-Smirnov statistic $K_S$. We use a mapping between simulation and physics units which minimises the mean value of $K_S$ obtained across all pairs of probes in all simulated cell lines. This leads to $\sigma =$ 17.6 nm.

To map the simulation time unit $\tau$ to a physical time we compare our simulations to *in vivo* measurements of chromatin dynamics. Specifically, we measure the polymer bead mean squared displacement as a function of lag time [MSD$(t)$] averaged over all beads in all simulations. We then use a mapping between simulation and physical units which gives the best fit to MSD$(t)$ curves obtained from Ref. 15, in which the authors performed particle tracking of probes at different chromatin regions in yeast. This leads to a value $\tau \approx 2.07 \times 10^{-3}$ s. The total run time of a simulation (after removing the first $7 \times 10^5\,\tau$ as detailed above) is therefore equivalent to about 27 minutes.

Energies are given in units of $k_B T$, and for simplicity we set the mass of all components (chromatin beads and proteins) to be the same ($m_i = m$ for all $i$). For computational efficiency we set $\gamma_i = 2m/\tau$; this is an approximation which means that the components have more inertia than in reality, but it renders the simulations much more efficient. This enables longer simulations to be performed such that we can assess the dynamics on relevant time scale (minutes).

As detailed above, we performed simulations of $L = 40,000$ bead long polymers containing 10 copies of the *Pax6* locus. For the majority of simulations we included 1000 proteins, and these switch back and forth between an active and inactive state at a rate $k_{\mathrm{sw}} = 10^{-3}\ \tau^{-1}$; using the above mapping this means proteins switch on a time scale of $0.48\,s$. In Figs. 4h-m and Supplementary Fig. 3a we present results from simulations using different values for these parameters as indicated in the relevant captions. Other important rates are related to loop extrusion and we used values: $k_{\mathrm{on}} = 2 \times 10^{-2}\,\tau^{-1}$, $k_{\mathrm{off}} = 2.5 \times 10^{-5}\,\tau^{-1}$, and $k_{\mathrm{ex}} = 2\,\mathrm{bp}/\tau$.

#### 2 Simulation Input Data

We use four sets of genomic data as an input to the simulations. Specifically, ATAC-seq data were used to identify active protein binding sites (we assumed all accessible DNA regions are binding sites); ChIP-on-chip data for H3K27ac were used to identify open chromatin regions; and ChIP-on-chip data for CTCF and the cohesin sub-unit Rad21 were used to identify loop anchor sites. We considered a 3 Mbp region around *Pax6* (chr2:104,000,000-107,000,000 mm9 genome build), mapping all features to beads representing 1 kbp of chromatin (i.e., this sets the resolution of the model at 1 kbp)

We previously obtained ATAC-seq data from each of the three cell lines as reported in Ref. 1 (available at GEO:GSE119656). As detailed in that reference, Nextera adaptors were trimmed from sequenced reads using the 'trim galore' utility, and aligned to the mm9 genome using bowtie2 [16]. Read pile-ups were generated and corrected for read depth using the 'genome coverage' tool from the Bedtools suite [17]. Peak calling was performed using MACS version 2.1.1 [18], with a $q$-value cut off of 0.05. In the model protein binding sites were identified by mapping the centre of each ATAC-seq peak to the corresponding 1 kbp polymer bead.

ChIP-on-chip data for H3K27ac, CTCF, and Rad21 were previously obtained from each of the three cell lines, again from Ref. 1 (available at GEO:GSE119659, GSE119658 and GSE120665). As detailed in that reference, the Bioconductor tool "Ringo" was used for pre-processing, normalisation, combining replicates and peak calling [19].

To identify "open chromatin" regions, we mapped the H3K27ac data to the set of polymer beads, and identified all beads which had an overlapping ChIP peak. For these beads we then did not include the additional springs which act to crumple the polymer.

To identify loop anchor sites we used the set of CTCF peaks which overlap with Rad21 peaks. We then found the directionality of each site by locating its underlying binding motif. The binding motif was obtained from the JASPAR database (matrix MA0139.1 [20]); the FIMO tool from the MEME suite [21] was used to search for and identify the direction of motifs within each peak. A given CTCF peak can contain multiple binding motifs, therefore a site's directionality was identified as the orientation of the motif with the highest score in terms of its match to the consensus motif. If there were motifs with similar score with different orientation we identified the peak as having both orientations. Peaks which did not overlap a CTCF motif were excluded. In each repeated simulation and copy of the locus we included only a subset of CTCF peaks, choosing them stochastically with a probability based on the peak height; this takes into account cell-to-cell variability in the occupancy of different CTCF sites. Finally, as for ATAC-seq peaks, we mapped the centre of the peak to a specific polymer bead.

#### 3 CaptureC data analysis

We previously generated CaptureC data for *Pax6* HIGH, ON and OFF cells [1] (data available at GEO:GSE120666). For this study we generated new CaptureC data for *Pax6* HIGH cells which had been treated with alpha amanitin or bleomycin as detailed in the Methods section in the main text. Oligonucleotides were designed to target restriction enzyme fragments at 13 locations across the locus as detailed in Supplementary Table 1.

For all CaptureC experiments we analysed the data using the capC-MAP software [22]. In brief, first reads were checked for adapter sequence and this was removed using the cutadapt tool [23]. An *in silico* restriction enzyme digestion was performed, breaking reads at the *DpnII* cutting sequence (GATC). The resulting read fragments were mapped to the mouse genome (mm9 build) using bowtie [24]. Aligned reads were filtered for PCR duplicates, and checked against the set of "target" restriction enzyme fragments for which oligonucleotide probes were designed. Only read pairs which identified a ligation event between a target and a (uniquely identifiable) "reporter" (i.e. non-target) fragment were retained. CaptureC interaction profiles were obtained for each target and sliding window binning and smoothing applied (using a window of 6 kbp and a bin width of 3 kbp; see Ref. 22 for details). Replicate data sets were compared, checked for consistence, and then combined to produce the final interaction profiles.

#### 4 Analysis of Simulation data

##### 4.1 Simulated CaptureC profiles

Simulated CaptureC interaction profiles were obtained from locus conformation generated from the HiP-HoP model following the same procedure as in Ref. 1. Specifically, we extract configurations from the simulations at $2000\,\tau$ intervals, and sample interactions stochastically. For a given viewpoint/target, for each copy of the locus in a simulation we select a bead at random from within the target region, and select a second bead at random from within the rest of that copy of the locus. We then accept this as an interaction with probability $P(x) = \exp(-x^2/x_0^2)$, where $x$ is the 3D separation of the two beads, and $x_0$ is a threshold set at $x_0 = 3.5\sigma$; this is repeated $L$ times where $L = 3000$ is the number of beads in that copy of the locus. We do this for each copy of the locus from a given cell type in each extracted configuration. We then repeat the whole procedure until the total number of accepted interactions is at least the same as than the number of reads typically obtained for a single viewpoint in our CaptureC data. From this procedure we generate a simulated interaction profile. Finally, we scale the profile such that the average number of reads per bead is the same in simulations and experiments.

##### 4.2 Mean squared displacement and mobility of simulated FISH probes

In order to calculate the mean squared displacement (MSD) for a given simulated FISH probe, we find the centre of mass of all beads covered by that probe in a given copy of the locus

$$\mathbf{r}_{\mathrm{CoM}} = \frac{1}{N}\sum_i \mathbf{r}_i,$$

where the sum runs over the $N$ beads covered by the probe. In this way we obtain the trajectory $\mathbf{r}_{\mathrm{CoM}}(t)$ and its MSD is calculated as

$$\mathrm{MSD}(t) = \langle [\mathbf{r}_{\mathrm{CoM}}(t+t_0) - \mathbf{r}_{\mathrm{CoM}}(t_0)]^2 \rangle,$$

where angle brackets denote an average over times $t_0$, and $t$ is the lag time. A MSD is obtained for each copy of the locus in each repeat simulation; in figures we plot the mean MSD over locus copies, and the standard error of this mean. Since we have 20 copies of the locus in each cell line (spread across 6 simulations), we have 20 independent MSDs, so the standard error is given by the standard deviation scaled by $1/\sqrt{20}$.

We define the mobility of a chromatin bead as the MSD at a fixed lag time $t^*$, specifically $\mathscr{M} = \mathrm{MSD}(t^*)$, and the mobility profile is obtained by calculating $\mathscr{M}$ for each polymer bead. We used a lag time $t^* = 10^4\,\tau$ (approximately equivalent to 20.7 s) which is short enough to give good statistics (i.e., small errors). We take the mean over $t_0$ and locus copies, and the error is given by the standard error in the mean (shown as a shaded region around the lines in Fig. 1E in the main text). We also considered mobility profiles using different lag times $t^* = 2\times 10^3\,\tau$ and $t^* = 2\times 10^4\,\tau$; while this shifted the mobility profile up or down, it did not significantly change its shape. A mobility can also be calculated for a probe covering multiple chromatin beads by using the MSD for the centre of mass of those beads (Supplementary Fig. 3c).

For an object freely diffusing in 3D, the mean squared displacement will grow linearly with time, $\mathrm{MSD}(t) = 6Dt$, where $D$ is the diffusion constant. More generally, $\mathrm{MSD}(t) \sim t^{\alpha}$; if the exponent $\alpha < 1$, the motion is said to be sub-diffusive. From the Rouse model for polymer dynamics[25] we expect the motion of a segment of a polymer to show an exponent $\alpha \approx 0.5$ (returning to diffusive behaviour at long lag times due to the polymer centre of mass motion). The inset in Fig. 2d reveals that the 40 kbp probe motion does show an exponent close to 0.5, at least up to lag times of about 250 s. A plateau in an MSD at long times is indicative of confined or constrained motion; given that our model includes several mechanisms which might constrain the motion we would expect a complicated MSD curve which goes through multiple regimes at long times.

##### 4.3 Local density and extruder density measure

To generate the local density profiles as shown in Fig. 2e, we calculated the density $\rho_i$ in the vicinity of chromatin bead $i$. This was defined as the total number of beads (polymer beads and proteins) within a sphere of radius $R^*$ centred at $\mathbf{r}_i$, divided by the sphere volume. We used $R^* = 3\sigma$, which is the same value used as the ‘contact threshold’ for generating simulated CaptureC profiles as detailed above. Finally, we average over the 20 copies of the locus for each cell line, and calculate the standard error in the mean (shown as a shaded region around the line in Fig. 2e in the main text; this is often of a similar thickness to the line).

We calculate the extruder density measure $\phi_e$ by counting, for chromatin bead $i$, the number of configurations in a simulation where there is an extruder spring at beads $i-3$ to $i+3$ (i.e. within a distance $3\sigma$ from bead $i$). This is divided by the total number of considered configurations. We then average over all 20 copies of the locus of a given cell type, and calculate a standard error in the mean. Profiles of $\phi_e$ across the locus are shown in Supplementary Fig. 4, with shaded regions around the lines showing the error. All $\phi_e$ values were used to generate the plot in Fig. 2i.

##### 4.4 Correlations between quantities

In Fig. 2g we show the relationship between the mobility and density of chromatin beads. Each point represents a single bead and data are included for all three cell lines, with the two quantities being calculated as detailed above. Similar scatter plots are shown in Supplementary Fig. 3a, but the same quantity is compared between two sets of simulations with different parameters.

In Figs. 2i we show the correlation between mobility and H3K27ac density. Here the data are given for 20 kbp (20 bead) windows rather than individual beads (we consider a window around each bead, from bead 10 to bead 2990 within each 3000 bead copy of the locus). The mobility value for a given window is taken to be the mean value over the 20 beads. The H3K27ac density is defined as the fraction of the 20 beads which have that property. In Fig. 2i we show the correlation between mobility and extruder density; here the values used are for each bead and are calculated as detailed above. In both panels data are used from all 20 copies of all three cell lines.

#### 4.5 Localness of interactions measure

To define a measure of the localness of interactions, for each chromatin bead we count interactions with other beads across the entire polymer. In this way, interactions with other locus copies are also included, meaning that these can be with chromatin regions further away than 3 Mbp (the size of the region around *Pax6* which we simulate). Since the surrounding chromatin environment in which the 3 Mbp locus is embedded in simulations is not representative of the *in vivo* environment, this measure should be considered only as a rough indication of interactions. Here, an interaction is defined as when two beads are closer together than $3.5\sigma$ in 3D space. We first exclude all interactions between beads separated by less than 5 kbp and more than 10 Mbp. Then the localness measure for a given bead within the locus is defined as the number of interactions with beads closer than (or equal to) 100 kbp away, divided by the number of interactions with beads further than 100 kbp away (Fig. 3a). This threshold distance is chosen such that interactions between promoters and distal enhancers will typically be counted as 'long ranged'. As with the other measures, we average over all copies of the locus for a given cell type. The error is the error in the mean, and in Fig. 3b this is shown as a shaded region around the lines (but is a similar size as the line thickness). In Fig. 3c values for mobility and localness for individual beads are used, and we include from all beads within all copies of the locus for all three cell types (beads in the 'spacer regions' between locus copies are not included).

#### 4.6 Dynamics of the locus structure

To characterise the locus structure we defined a vector

$$\mathbf{X} = (x_{\mathrm{UP}}, x_{\mathrm{UD}}, x_{\mathrm{PD}}),$$

where $x_{\mathrm{UP}}$, $x_{\mathrm{UD}}$, and $x_{\mathrm{PD}}$ are the separations of the centre of mass of the URR and *Pax6*, URR and DRR, and *Pax6* and DRR FISH probe regions respectively. In this way $\mathbf{X}$ is a vector in a three-dimensional configuration space, and we can track its trajectory in time $\mathbf{X}(t)$ (Figs. 4a-e). The $\mathbf{X}$ vector can also be obtained for a single fixed cell in 3-colour FISH experiments.

The 'shape-change' parameter is then defined as the mean squared displacement of $\mathbf{X}(t)$

$$S(t) = \langle (\mathbf{X}(t+t_0) - \mathbf{X}(t_0))^2 \rangle,$$

where $t$ is the lag time, and the angled brackets denote an average over trajectories and times $t_0$ in the same way as the mean squared displacement calculation detailed above.

A measure of the mean size of the locus at time $t$ is given by the root mean squared of the vector $\sqrt{\langle \mathbf{X}(t) \rangle}$, where the angle brackets denote an average over time points and repeat simulations of the locus (Supplementary Fig. 2d left shows the distribution of locus size over all configurations in all copies of the locus for a given cell type). This same measure can be obtained for a single fixed cell from FISH experiments (distributions are show in Supplementary Fig. 2c left).

A measure of the variability of the locus configuration can be obtained from the volume of cloud of points in configuration space (Fig. 4d, all points along all of the trajectories from each copy of the locus for a given cell line are included). To estimate this we calculate the radius of gyration of of the points; this leads to

$$\text{variability} = \left( \frac{1}{NM} \sum (\mathbf{X} - \langle \mathbf{X} \rangle)^2 \right)^{3/2},$$

where the sum runs over the $N$ points obtained from each of $M$ repeat simulations, and the $3/2$ power ensures units of volume. The same measure can be made using $\mathbf{X}$ vectors obtained from single fixed cells from FISH experiments (Supplementary Fig. 2c right).

#### 4.7 Collision interval and interaction duration

In Fig. 5b,c we consider the collision interval and interaction duration for interactions between the *Pax6* promoters and the two enhancers (URR and DRR). For each of these elements there were several ATAC-seq peaks in close proximity, so we considered a region at each element which covered all of the peaks. Specifically at the URR we considered a 13 chromatin bead region (chr2:105,444,500-105,457,500), at *Pax6* we considered an 11 bead region (chr2:105,509,500-105,520,500), and the the DRR an 11 bead region (chr2:105,629,500-105,640,500 all coordinates mm9 reference genome). This threshold was chosen as it is roughly equal to the radius of gyration of one of the 11 bead regions. From a given trajectory, the duration of periods where two elements are together were recorded as 'interaction durations', and the duration of periods where two elements are apart were recorded as 'collision intervals'. In plots the mean times for these events across all 20 trajectories for each cell line is shown. Polymer configurations were only retained at intervals of $2 \times 10^3 \tau$, so this limits the accuracy of these measurements. In Fig. 5d we consider 10 kbp regions which tile the locus between *Rcn1* and *Elp4*, and perform the same mean collision interval and interaction duration measurements for every possible pair of regions in order to build up a map.

#### 5 Two models for simulating alpha amanitin treatment

As detailed in the main text, we considered two alternative scenarios for including alpha amanitin treatment in simulations.

In Model 1 (Fig. 5c) all proteins were removed from the system, as these represent complexes of transcription factors and RNA polymerase. The simulation was then run for $1.5 \times 10^6 \tau$, with loop extrusion continuing. Only configurations from the final $8 \times 10^5 \tau$ of the simulation were used to generate plots (Figs. 5d,g-i) to allow for a relaxation after the simulated 'treatment'. Six independent simulations, with a total of 20 copies of each locus for each cell line were performed.

In Model 2 (Fig. 5e) protein-chromatin bonds were instead made permanent at the point of treatment. We first ran a standard simulation for $1.5 \times 10^6 \tau$. Then to simulate the treatment we identify all proteins which are within a distance $1.8\sigma$ of a chromatin bead; a new harmonic bond was then added between these protein and the chromatin beads

$$U_{AA}(r_{i,j}) = U_{WCA}(r_{i,j}) + K_{AA}(r_{i,j} - r_0)^2$$

with $K_{AA} = 20k_BT/\sigma^2$ and $r_0 = 1.1.\sigma$. At the same time, all other protein-chromatin attractions were switched off. The simulation was then continued for a further $1.5 \times 10^6 \tau$, with loop extrusion continuing. Again, only configurations from the final $8 \times 10^5 \tau$ of the simulation were used to generate plots (Figs. 5f-i) to allow for a relaxation after the simulated treatment.
